# Alterations of specific inhibitory circuits of the prefrontal cortex underlie abnormal network activity in a mouse model of Down syndrome

**DOI:** 10.1101/2020.02.24.963041

**Authors:** Javier Zorrilla de San Martin, Cristina Donato, Jérémy Peixoto, Andrea Aguirre, Vikash Choudhary, Angela Michela De Stasi, Joana Lourenço, Marie-Claude Potier, Alberto Bacci

## Abstract

Down syndrome (DS) results in various degrees of cognitive deficits. In DS mouse models, recovery of behavioral and neurophysiological deficits using GABA_A_R antagonists led to hypothesize an excessive activity of inhibitory circuits in this condition. Nonetheless, whether over-inhibition is present in DS and whether this is due to specific alterations of distinct GABAergic circuits is unknown. In the prefrontal cortex of Ts65Dn mice (a well-established DS model), we found that the dendritic synaptic inhibitory loop formed by somatostatin-positive Martinotti cells (MCs) and pyramidal neurons (PNs) was strongly enhanced, with no alteration of their excitability. Conversely, perisomatic inhibition from parvalbumin-positive (PV) interneurons was unaltered, but PV cells of DS mice lost their classical fast-spiking phenotype and exhibited increased excitability. These microcircuit alterations resulted in reduced pyramidal-neuron firing and increased phase locking to cognitive-relevant network oscillations *in vivo*. These results define important synaptic and circuit mechanisms underlying of cognitive dysfunctions in DS.

## Introduction

Down syndrome (DS) is a condition caused by full or partial trisomy of human chromosome 21, characterized by various physical and neurological features including mild to severe intellectual disability (Antonarakis et al., 2020). Individuals with DS present important deficits in cognitive tasks known to depend on the anatomical and functional integrity of the frontal lobe (Lee et al., 2015). Moreover, DS is also associated with other CNS-mediated phenotypes, including an ultra-high risk for developing Alzheimer’s disease and high rates of autism for which the mechanisms are unknown (DiGuiseppi et al., 2010; Wiseman et al., 2015). Interventions to ameliorate DS-mediated cognitive dysfunctions are limited. The development of interventions for this vulnerable group of individuals can be achieved through a better understanding of the mechanisms underlying core feature of DS, such as intellectual disability.

Importantly, mouse models of DS recapitulate several cognitive deficits of this condition (Herault et al., 2017; Olmos-Serrano et al., 2016). One of the best-characterized mouse models is the Ts65Dn mice (herein referred to as Ts), which carry a partial trisomy of a segment of the mouse chromosome 16 (Davisson et al., 1993) and recapitulates several dysfunctions present in DS individuals, such as reduced birthweight, male sterility, abnormal facial appearance and several cognitive impairments, including executive functions, such as working memory and cognitive flexibility (Olmos-Serrano et al., 2016).

Executive functions depend on the integrity of the prefrontal cortex (PFC), which is proposed to play an essential role in the synchronization of task-relevant, large-scale neuronal activity (Helfrich and Knight, 2016). An important network correlate of this synchronization is represented by neuronal oscillations: rhythmic fluctuations of the electrical activity of single neurons, local neuronal populations and multiple neuronal assemblies, distributed across different brain regions (Buzsáki and Wang, 2012). Oscillations are the result of a balanced and coordinated activity of excitatory pyramidal neurons (PNs) and a rich diversity of inhibitory neurons that use γ-aminobutiric acid (GABA) as neurotransmitter. In particular, parvalbumin (PV)-positive inhibitory interneurons form synapses onto the perisomatic region of PNs, thus tightly controlling their spiking activity and driving fast network oscillations in the γ-frequency range (30–100 Hz) (Buzsáki and Wang, 2012). γ-Oscillations are necessary for several PFC cognitive functions, such as sustained attention (H. Kim et al., 2016) and cognitive flexibility (Cho et al., 2015). Conversely, Martinotti cells (MCs) are somatostatin-positive interneurons that inhibit distal dendrites of PNs, thereby controlling the integration of distal dendritic glutamatergic synaptic inputs originating from different regions of the brain (Tremblay et al., 2016). Dendritic integration of multi-pathway inputs is necessary for working memory (Abbas et al., 2018; D. Kim et al., 2016). Therefore, PV interneurons and MCs represent two major cortical inhibitory circuits, characterized by a precise division of labor during cortical activity. Both forms of inhibition were shown to be involved in the entrainment of network oscillations (Cardin et al., 2009; Sohal et al., 2009; Veit et al., 2017) and in the cognitive performance during medial (m)PFC-dependent tasks (Abbas et al., 2018; Cho et al., 2015; Clem and Cummings, 2020). In particular, inhibition from SST interneurons plays a crucial role in mPFC-dependent memory (Abbas et al., 2018; Clem and Cummings, 2020).

Recovery of behavioral and neurophysiological deficits underlying cognitive impairments using GABA_A_ receptor blockers led to hypothesize that intellectual deficits in DS are produced by an excessive activity of inhibitory circuits (Fernandez et al., 2007; Zorrilla de San Martin et al., 2018). Nonetheless, direct evidence for over-inhibition in DS is lacking. Moreover, given the anatomical, molecular and functional diversity of cortical inhibitory neurons (Tremblay et al., 2016), the functional implications of this hypothesis at the network level, as well as the involvement of specific GABAergic circuits remain obscure.

Interestingly, whereas broad-spectrum GABA_A_R antagonists are not clinically viable (as they can yield undesired seizure-like activity and/or anxiety), treatment of Ts mice with selective and partial negative allosteric modulators of α5-containing GABA_A_Rs (α5 inverse agonist or α5-IA) reverse cognitive behavioral and long-term synaptic plasticity deficits in DS mice (Braudeau et al., 2011; Duchon et al., 2019; Martínez-Cué et al., 2013; Schulz et al., 2019). Importantly, neocortical dendritic synaptic inhibition of PNs from MCs relies on α5-containing GABA_A_Rs (Ali and Thomson, 2008). The preference for this specific GABA_A_R subunit was also recently demonstrated at the equivalent hippocampal dendritic inhibitory circuit (Schulz et al., 2019, 2018), raising the question of whether dendritic inhibition is specifically altered in DS.

Here we found that the dendritic synaptic inhibitory loop formed by MCs and PNs was strongly potentiated in Ts mice, with no alteration of either cell-type excitability. Conversely, the perisomatic synaptic inhibitory loop from PV cells onto PN cell bodies was unaffected in Ts mice. Strikingly, however, PV-cell excitability was strongly altered: these interneurons did not display their typical fast-spiking behavior and exhibited enhanced excitability. At the network level *in vivo*, these inhibitory microcircuit-specific alterations resulted in significant reduction of putative PN firing, which in turn was more tuned to β- and low γ-oscillations (20-40 Hz). These results confirm over-inhibition in DS, and reveal unexpected functional alterations of specific GABAergic circuits in this condition.

## Results

### Synaptic enhancement of dendritic inhibition in DS

Cognitive and synaptic plasticity deficits in Ts mice are recovered by systemic application of a selective negative allosteric modulator of α5-containing GABA_A_Rs, α5IA (Braudeau et al., 2011; Duchon et al., 2019; Martínez-Cué et al., 2013; Schulz et al., 2019). α5-GABA_A_Rs are expressed at synapses between dendrite-targeting interneurons: MCs and O-LM in the neocortex (Ali and Thomson, 2008) and hippocampus (Schulz et al., 2018), respectively. We therefore tested whether dendritic inhibition of PNs by MCs could be affected in Ts mice.

We crossed Ts65Dn with GFP-X98 mice, which in the barrel cortex were shown to bias GFP expression in MCs (Ma, 2006). Accordingly, in the mPFC of these mice, GFP was expressed by a subset of SST-positive interneurons (Figure 1 – figure supplement 1a), exhibiting a widely branched axonal plexus in L1, characteristic of dendrite-targeting inhibitory MCs (Figure 1 – figure supplement 1b,c). Using dual whole-cell patch-clamp in acute mPFC slices, we recorded unitary inhibitory postsynaptic currents (uIPSCs) in MC-PN connected pairs (Figure 1a) and found that dendritic MC-PN synaptic inhibition relied on α5-containing GABA_A_Rs in both Ts and Eu mice. Indeed, bath application of the selective negative allosteric modulator of α5-containing GABA_A_Rs, α5IA (100 nM; (Sternfeld et al., 2004), produced a significant reduction of uIPSCs that was close to the maximal potency of the drug (~40%; Dawson et al., 2006) in both genotypes (Figure 1b, Supplementary Table 1). Interestingly, MC-mediated uIPSC amplitudes were significantly larger and failure rate significantly smaller in Ts compared to Eu. Moreover, the total amount of charge (Q) transferred during a train of 5 action potentials was near 5-fold larger in Ts than in Eu (Figure 1c, Supplementary Table 1). In addition, glutamatergic recruitment of MCs by PNs was also stronger in Ts than Eu mice. Unitary excitatory postsynaptic currents (uEPSCs) in connected PN-MC pairs exhibited larger amplitude, lower failure rate and larger charge transfer in Ts than Eu mice (Figure 1d, Supplementary Table 2).

**Figure 1:**
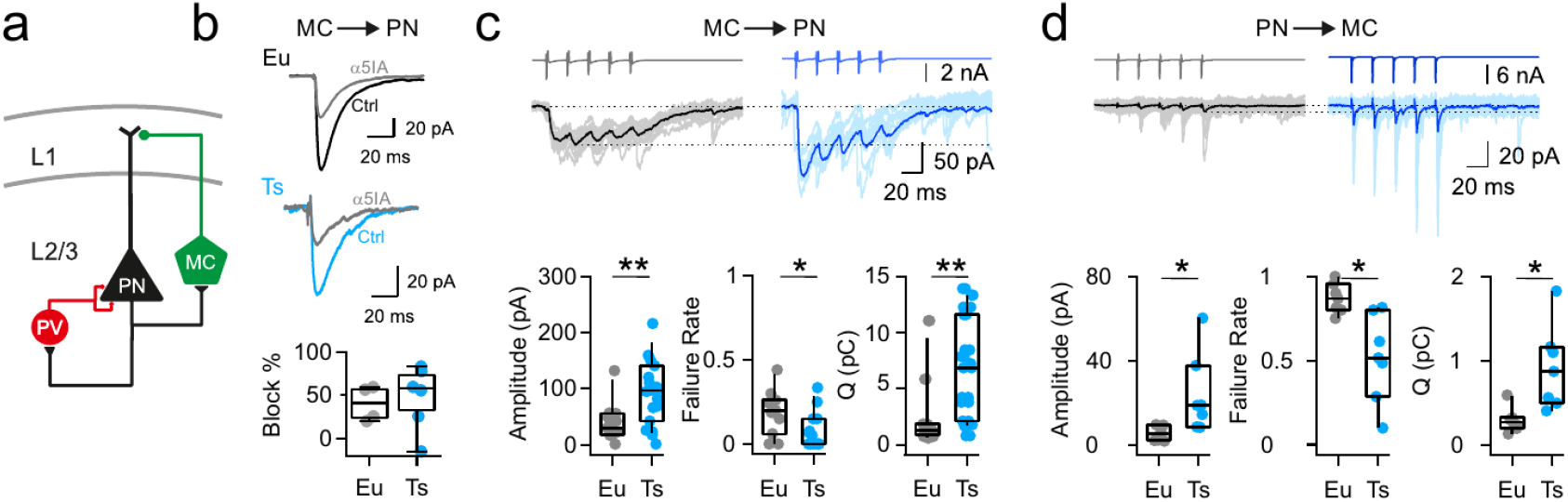
Synaptic enhancement of dendritic inhibition in DS. **a,** Schematic of cortical inhibitory circuits involving PV interneurons, MCs and PNs. **b,** Representative voltage-clamp current traces of uIPSCs recorded in MC-PN connected pair in a Eu (top) and a Ts (bottom) mouse, before (Ctrl) and after (α5IA) application of the α5-GABA_A_R inverse agonist α5IA (100 nM). Shown are averages of 50 traces Right, bottom: population data of uIPSC amplitude block by α5IA in Eu and Ts mice. **c,** Top: Representative traces of uIPSCs (lower traces) elicited by a train of 5 presynaptic action currents (50 Hz) in the MC (upper traces). Eu: individual (gray) and average (black) traces; Ts: individual (light blue) and average (blue) traces. Bottom panels: population data of uIPSC amplitude (p=0.0094, Mann-Withney U-test), failure rate (p=0.0201, Mann-Withney U-test) and total charge transferred during the 5 APs train (Q, p=0.006, Mann-Withney U-test; n=11 and 15 pairs for Eu and Ts respectively). **d,** Same as in c, but for glutamatergic uEPSCs triggered by action currents in presynaptic PNs and recorded in postsynaptic MCs (Amplitude: p=0.0398, Mann-Withney U-test; Failure Rate: p=0.0107, Mann-Withney U-test; Q: p=0.0157, Students T-test; n=7 for both, Eu and Ts).

Importantly, in Ts mice, the short-term dynamics of both MC-PN GABAergic and PN-MC glutamatergic synaptic transmission were unaltered. Indeed, inhibitory dendritic inhibition of PNs exhibited short-term depression in both genotypes. Likewise, glutamatergic recruitment of MCs displayed the classical strong facilitating characteristics (Silberberg and Markram, 2007) in both Ts and Eu mice.

Altogether, these results indicate that the dendritic inhibitory loop involving MCs and PNs is strengthened in Ts mice. Both output GABAergic synapses from MCs and their recruitment by local glutamatergic synapses were stronger and more reliable than in DS mice, as compared to their euploid littermates.

### Unaltered excitability and morphology of MCs and PNs in Ts mice

Alteration of the MC-PN-MC synaptic loop can be associated to changes in intrinsic excitability and morphological features. We therefore tested whether passive properties, single action potentials and firing dynamics were altered in both PNs and MCs. In addition, we filled neurons with biocytin and we quantified their dendritic and axonal arborizations. Input-output spiking activity of both PNs and MCs, was assessed by injecting increasing depolarizing 1 s-long currents. The firing frequency vs. injected current (*f-i*) curve was similar in both cell types in Eu and Ts mice (Figure 2a,b; Supplementary Table 3–4). Furthermore, single-action potential features (such as spike threshold, peak and duration), and passive properties (such as input resistance, resting membrane potential and time constant) were similar in both genotypes (Figure 2 – figure supplements 1 and 2; Supplementary Tables 5–6). Importantly, the density of GFP-expressing MCs, and, in general, of SST-positive interneurons was similar in both genotypes (Figure 2c; Figure 2 – figure supplement 3).

**Figure 2:**
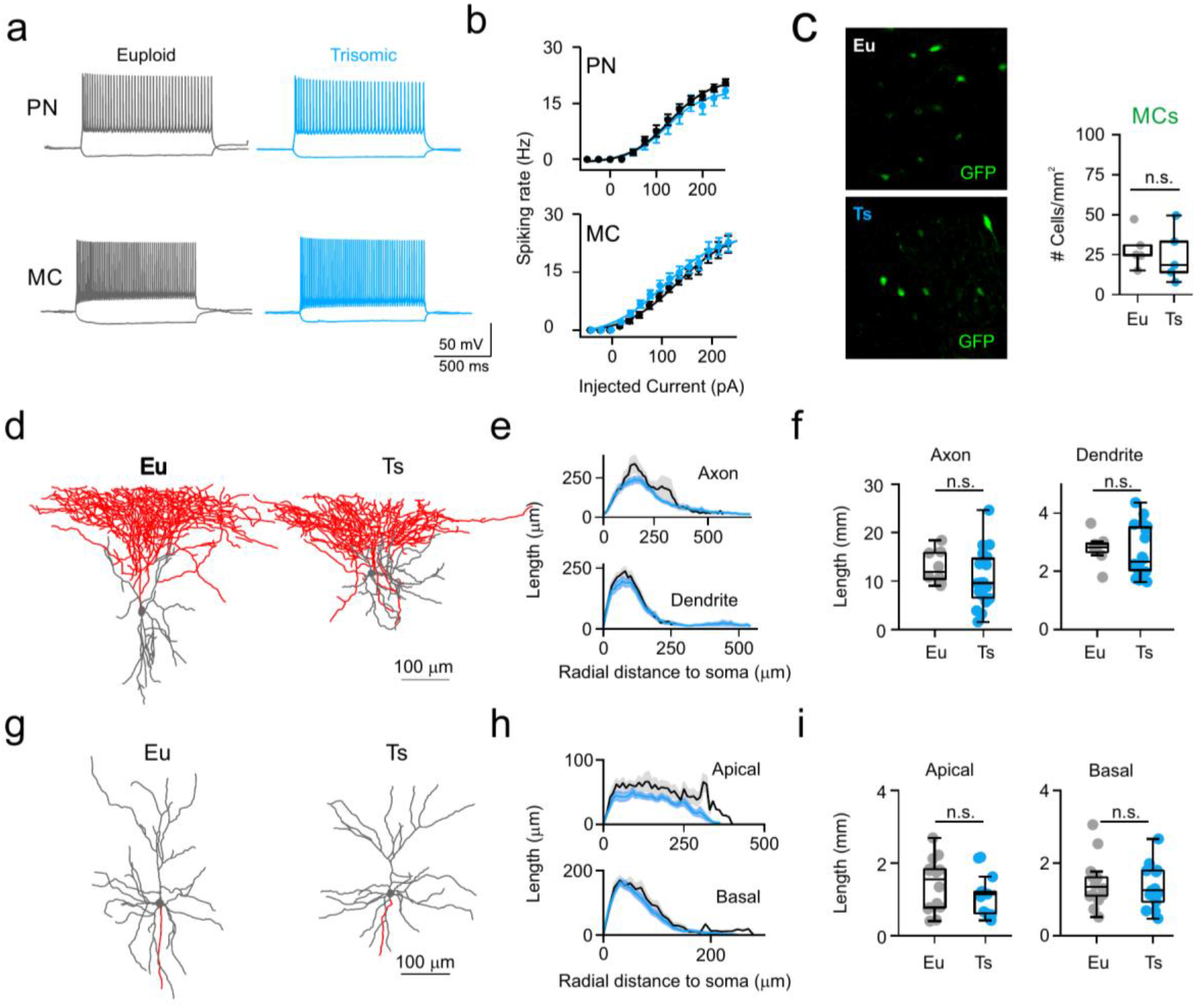
Normal excitability and morphology of PNs and MCs in Ts mice. **a,** Representative current-clamp traces of membrane potential responses to injections of current steps of increasing amplitude applied to PNs (above) and MCs (below) from Eu (gray) and Ts (blue) mice. **b,** Spiking frequency as function of injected current. Population data from PNs (left, genotype factor F_(1, 598)_=3.444, p=0.064, two-way ANOVA; n=25 and 23 cells for Eu and Ts correspondingly) and MCs (right, genotype factor F_(1, 660)_=0.004960, p=0.9439, two-way ANOVA; n=26 and 20 cells for Eu and Ts correspondingly). **c,** Count of MC somas in the mPFC of Eu and Ts mice. Left: epifluorescence images of immuno-labeled GFP-expressing MCs in Ts::X98 coronal slices. Right: population data for both Eu and Ts. **d,** Representative reconstruction of biocytin filled L2/3 MCs from Eu and Ts mice. Gray: somatodendritic region, red: axon. **e,** Scholl analysis of MC axonal and dendritic length between concentric circles of increasing radial steps of 10 μm. **f,** Population data of total axonal (left) and dendritic (right) length. **e,f,** n=18 and 8 neurons for Eu and Ts respectively. **g,** Representative reconstruction of biocytin filled PNs from Eu and Ts mice. Gray: somatodendritic region (including apical and basal dendrites), red: axon. **h-i,** same as in e-f, but for apical and basal dendrites of PNs (F _(1, 25)_=2.487, p=0.1273 2-way ANOVA for apical dendrites; F _(1, 25)_= 0.3521, p=0.5583 2-way ANOVA for basal dendrites). **h,i,** n=14 and 13 neurons for Eu and Ts respectively. **c,f,i,** Boxplots represent median, percentiles 25 and 75 and whiskers are percentiles 5 and 95. Points represent values from individual synapses (b,c), mice (f), neurons (i,l). *: p < 0.05; **: p < 0.01

Increased MC-PN GABAergic transmission synaptic transmission in Ts mice can be attributed to axonal sprouting of MCs and/or increased dendritic branching of PNs. We performed a morphometric analysis of both cell types and found that the spatial distribution and total length of axons and dendrites of MCs were similar in both genotypes (Figure 2d-f). Likewise, both apical and basal dendrite arborizations of PNs were indistinguishable in Eu and Ts mice (Figure 2g-i).

These experiments and those illustrated in Figure 1 indicate that the increased dendritic inhibitory loop involving MCs and PNs can be specifically attributable to alteration of synapses between these two cell types.

### Excitability of PV cells, and not their perisomatic control of PNs, is strongly altered in Ts mice

Is the synaptic enhancement of the dendritic inhibitory loop involving MCs a specific alteration or a general feature of glutamatergic and GABAergic synapses in Ts mice? To address this question, we measured glutamatergic recruitment onto, and synaptic inhibition from, another prominent interneuron class, PV basket cells. These interneurons are characterized by their ability of firing at high frequencies, non-adapting trains of fast action potentials. These properties, along with the perisomatic inhibitory basket formed by their axon around the soma of PNs, make PV interneurons an efficient regulator of PN output. We thus crossed Ts65Dn with PValb-tdTomato mice, a line that expresses TdTomato specifically in PV-positive interneurons (Kaiser et al., 2016). We recorded uIPSCs and uEPSCs (Figure 3a-d) from pairs of synaptically connected PNs and PV cells. The amplitude, failure rate and charge transfer of trains of uIPSCs evoked by action potentials in presynaptic PV-INs were similar in both genotypes (Figure 3a,b; Supplementary Table 7). Likewise, amplitudes, failure rates and charge transfer of uEPSCs elicited by PN firing was indistinguishable in Eu and Ts mice (Figure 3c,d; Supplementary Table 8).

**Figure 3:**
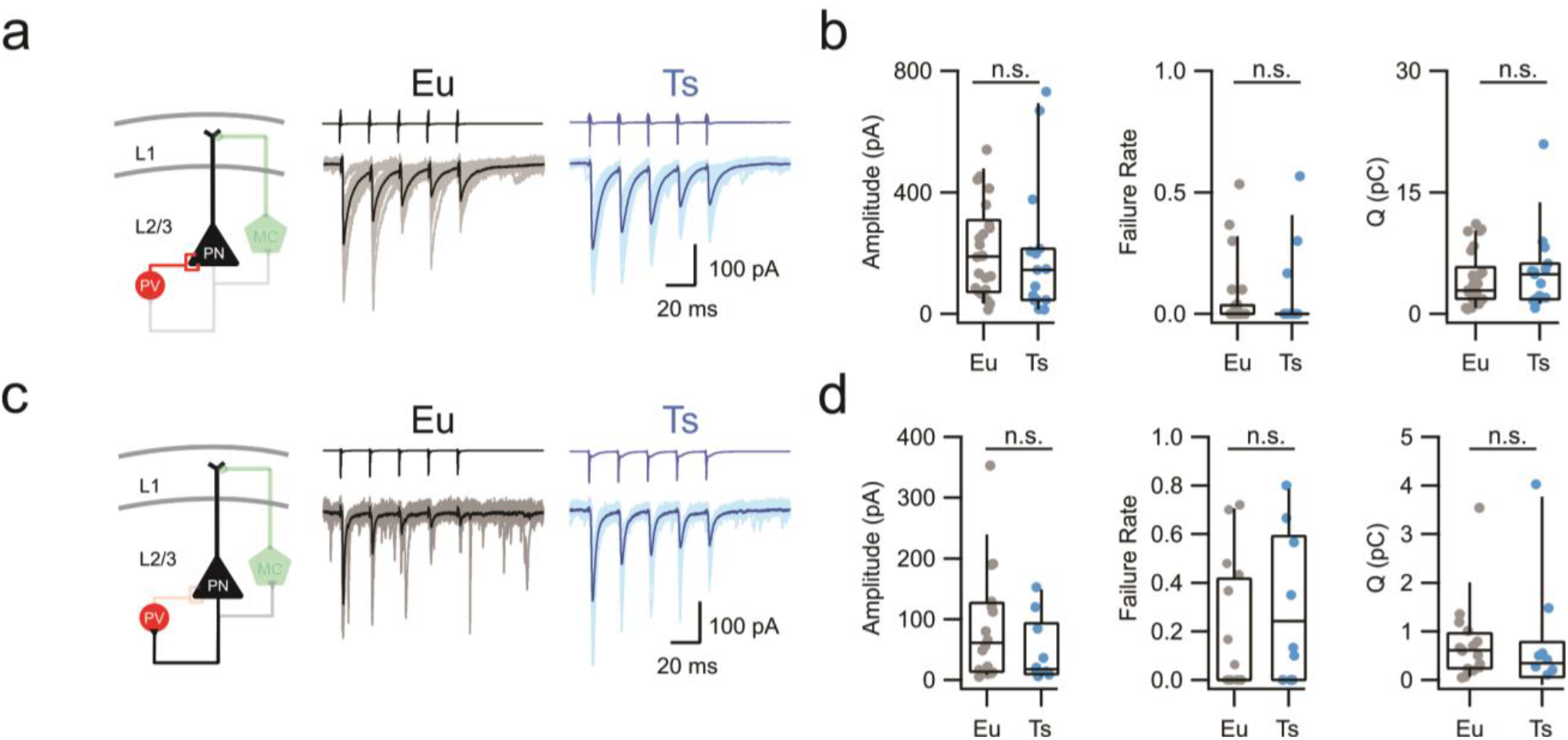
Perisomatic inhibition by PV-INs is normal in Ts mice. **a,** Left: scheme of the PV-PN perisomatic inhibitory circuit assessed using dual whole-cell patch-clamp recordings. Right: representative voltage-clamp traces corresponding to uIPSCs evoked upon application of 5 action currents (50 Hz) to the presynaptic PV-IN. Eu: individual (gray) and average (black); Ts: individual (light blue) and average (blue) traces are superimposed. **b,** Population data of uIPSC amplitude (p=0.4901, Mann-Withney U-test), failure rate (p=0.7185, Mann-Withney U-test) and total charge transferred during the 5 APs train (Q, p=0.579, Mann-Withney U-test, n=26 and 15 pairs for Eu and Ts correspondingly). **c-d,** Same as in a-b, but for glutamatergic uEPSCs triggered in the presynaptic PN and recorded in the postsynaptic PV IN in both genotypes (Amplitude: p=0.233, Mann-Withney U-test; Failure Rate: p=0.214, Mann-Withney U-test; Q: p=0. 3711, Mann-Withney U-test; n=15 and 9 pairs for Eu and Ts correspondingly).

Surprisingly, however, when we tested intrinsic excitability of PV cells, we found that, in Ts mice, these interneurons required one third less current to fire action potentials, as compared to their Eu littermates. (Figure 4a-c). Moreover, PV cells in Ts mice could not sustain high-frequency firing in response to 2 s-long depolarization, and maximal spike rate was thus near half of that reached by PV cells in Eu mice (Figure 4a-c; Supplementary Table 9). Notably, in Ts mice, action potential width was 1.7-fold wider and input resistance was 1.9-fold of that observed in Eu mice (Figure 4d). Conversely, action potential threshold and amplitude were not affected (Figure 4 – supplement figure 1; Table 10). Similarly to SST cells, the density of PV-INs in mPFC was similar in the two genotypes (Figure 4f).

**Figure 4:**
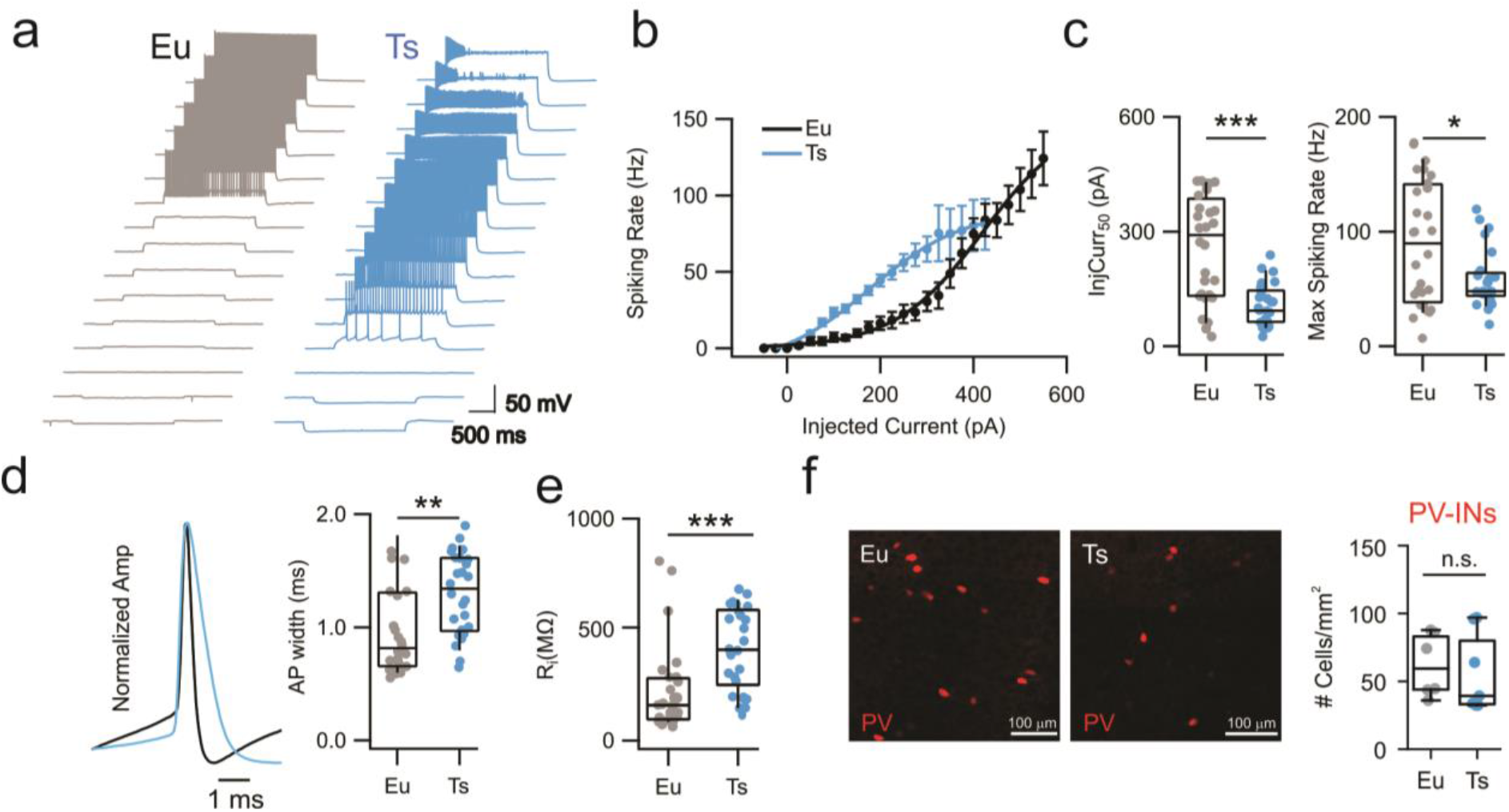
Altered excitability of PV cells in Ts mice. **a,** Representative voltage traces in response to current steps of increased amplitudes applied to to PV-INs from Eu (gray) and Ts (blue). **b,** Spiking rate as function of injected current Eu (black) and Ts (blue) mice (Genotype factor F_(1, 1040)_=53.90, P<0.0001, two-way ANOVA; n=28 and 26 cells for Eu and Ts correspondingly). **c,** Left: Population data of the injected current intensity required to reach 50% of maximal spiking rate estimated for each cell in Eu and Ts mice (p=4.55e^-6^, Mann-Withney U-test). Right: Maximal spiking rate reached upon current injection (p=0.04, Mann-Withney U-test, n=28 and 26 cells for Eu and Ts correspondingly). **d,** Left: representative action-potential traces, scaled to the peak, from Eu (black) and Ts (blue) mice. Right: population data of AP width in the two genotypes. **e,** Population data of input resistance measured in Eu and Ts PV-INs. **f,** Quantification of PV-INs somas in the mPFC of Eu and Ts mice. Left: Epifluorescence images of immunolabeled PV cells in coronal slices from Ts::PV mice. Right: density of PV cells Eu and Ts (n=6 and 7 mice for Eu and Ts respectively; p=0.2602, Mann-Withney test). *: p < 0.05; **: p < 0.01

Altogether, these results indicate that, contrary to dendritic inhibition, the synaptic efficiency of the perisomatic feedback loop mediated by PV-INs was normal in Ts mice. Yet, PV-INs of Ts mice lost their characteristic electrophysiological fast-spiking signature, and their excitability was dramatically increased.

### Reduced spiking activity *in vivo* and increased tuning with network oscillations in Ts65Dn mPFC

Increased dendritic inhibition by the MC-PN loop, altered spiking activity of PV cells and their increased excitability will likely strongly influence the spiking properties and dynamics of mPFC PNs in vivo during spontaneous network activity. In order to assess the activity of the mPFC *in vivo*, we performed simultaneous local field potential (LFP) and loose-patch, juxtacellular recordings from layer 2/3 putative PNs to monitor their spiking dynamics related to overall network activity (Figure 5a). *In vivo* recordings exhibited typical oscillatory activity consisting of UP and DOWN states (Ruiz-Mejias et al., 2011). UP and DOWN states were similar in frequency and duration in Ts mice and their Eu littermates (Figure 5 – figure supplement 1). UP states were enriched in γ-band activity (30-100 Hz) and exhibited increased probability of spiking activity (Ruiz-Mejias et al., 2011) (Figure 5a). Juxtacellular recordings from individual mPFC putative PNs revealed a near 50% decrease in the overall spiking rate in Ts mice as compared to their Eu littermates (Figure 5b, Supplementary Table 11). Analysis of LFP power spectral density (PSD) did not show a strong difference in the two genotypes (Figure 5c). Interestingly, however, when we analyzed LFP waveform specifically around periods of neuronal spiking activity (spike-triggered LFP or stLFP), we found that, in both genotypes, average stLFPs exhibited marked voltage deflections, indicating that spike probability was not randomly distributed but locked to LFP oscillations (Figure 5d). The peak-to-peak amplitude of the stLFP was much larger in Ts than in Eu mice (Figure 5d,e, Supplementary Table 11), and, remarkably, the spectral power of the stLFP was largely increased in Ts mice, selectively in the β-γ-frequency band (Figure 5f). In order to quantitatively assess whether PN spikes were differently locked to the phase of network oscillations, we measured the pairwise phase consistency (PPC), which is an unbiased parameter to determine the degree of tuning of single-neuron firing to network rhythmic activity of specific frequencies (Perrenoud et al., 2016; Veit et al., 2017). Ts mice exhibited significantly higher PPC values than Eu littermates for frequency bands ranging between 10 and 60 Hz (Figure 4g, Supplementary Table 12), thus revealing stronger phase locking selectively with β- and low γ-oscillations.

**Figure 5:**
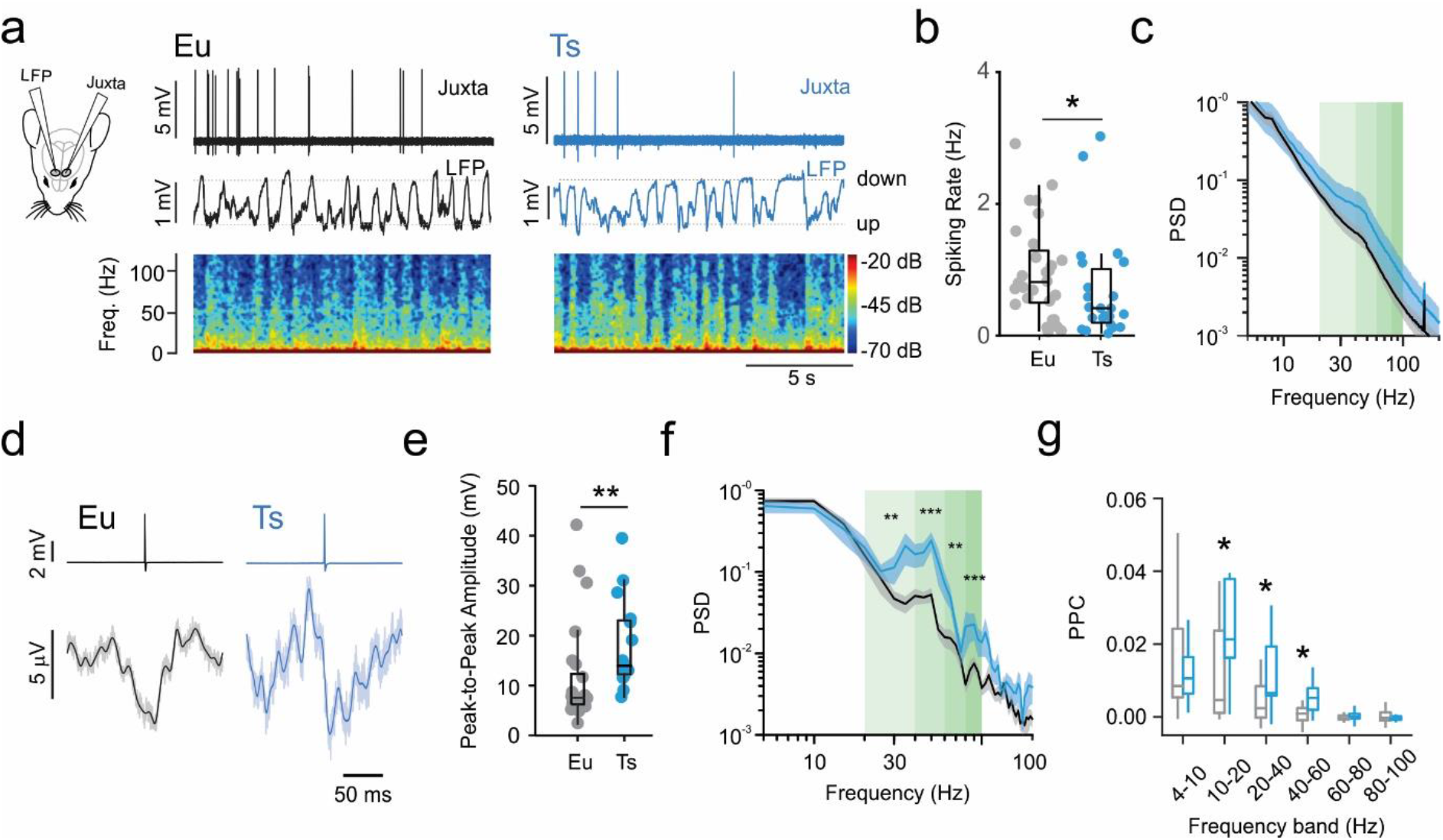
Reduced spiking activity *in vivo* and increased tuning with network oscillations in Ts65Dn mPFC. **a,** Left: scheme depicting simultaneous local field potential (LFP) and juxtacellular patch clamp recordings in the mPFC. Right: representative juxtacellular (top trace), LFP (middle trace) and spectrogram (bottom) recorded in Eu (black traces) and Ts (blue traces) mice. **b,** Average spiking rate from individual cells. Population data (n=28 cells, 5 mice and 23 cells, 6 mice for Eu and Ts respectively; p=0.03, Mann-Withney U-test). **c,** Power Spectral Density (PSD) from Eu and Ts mice. Shaded green areas correspond to β-γ-frequency ranges. **d,** Representative portions of averaged LFP around aligned spike (spike-triggered LFP or stLFP). Top: Average traces of aligned spikes from a putative layer 2/3 PN recorded in a Eu (black) and Ts (blue) mouse. Bottom: average of the corresponding stLFPs. Light line: raw averaged trace, thick dark line: low pass filtered trace (cutoff: 100 Hz). **e,** Average plots of stLFP peak-to-peak amplitude in Eu and Ts mice. **f,** Power spectral density of stLFPs. Shaded green areas correspond to β-γ-frequency ranges. **g,** Pairwise Phase Consistency (PPC) calculated for specific frequency bands (4-10Hz: p=0.427, 10-20Hz: p=0.02145, 20-40Hz: p=0.0418, 40-60Hz: p=0.01638, 60-80Hz: p=0.2308, 80-100Hz: p=0.3161, Mann-Withney U-test). e-g,: n=23 cells, 5 mice and 11 cells, 6 mice for Eu and Ts respectively. Boxplots represent median, percentiles 25 and 75 and whiskers are percentiles 5 and 95. *: p < 0.05; **: p < 0.01.

Altogether, these results indicate that mPFC PNs in Ts mice fire less than their Eu littermates, but their spontaneous spiking activity is more strongly tuned to cognition-relevant network activity. Both reduced firing rate and increased phase locking with fast oscillations are consistent with increased activity of inhibitory interneurons (Cardin et al., 2009; Sohal et al., 2009; Veit et al., 2017).

## Discussion

In this study, we analyzed synaptic and intrinsic properties of two major inhibitory circuits of the prefrontal cortex in a relevant mouse model of DS. We found specific alterations of distinct GABAergic circuits. In particular, we demonstrate that the dendritic inhibitory synaptic loop involving MCs and PNs was strongly potentiated in Ts, as compared to euploid control mice. In contrast, the synaptic perisomatic inhibitory control of PNs by PV cells was not affected in Ts mice. Strikingly, however, the excitability of PV cells was profoundly altered. These GABAergic circuit-specific alterations well explain the reduced spike rate and coupling with network activity of PNs of Ts mice in vivo.

We analyzed MCs, which are a subset of SST interneurons, specialized in inhibiting the distal portion of apical dendrites of PNs, by ascending their axons in L1 (Kawaguchi and Kubota, 1998), where PN dendrites integrate top-down input originating from distal brain areas. In the hippocampus, dendritic inhibition strongly regulates PN dendrite electrogenesis and supra-linearity (Lovett-Barron et al., 2012), likely modulating the emergence of burst firing (Royer et al., 2012). In the neocortex, integration of top-down information in L1 and consequent dendrite-dependent generation of bursts was hypothesized to underlie the encoding of context-rich, salient information (Larkum, 2013). Our results indicate a specific synaptic strengthening of the dendritic inhibitory loop involving MCs in Ts mice, suggesting a major impact on PN dendritic integration and electrogenesis. Enhanced dendritic inhibition by MCs in DS could underlie the deficits of long-term plasticity of glutamatergic synapses similar to those observed in the hippocampus of Ts mice. Indeed, these LTP deficits can be recovered by treatment of allosteric modulators of α5-GABA_A_Rs (Fernandez et al., 2007; Kleschevnikov et al., 2004; Schulz et al., 2019). Likewise, the enhanced MC-PN inhibitory loop in Ts mice shown here can provide a mechanism for the rescue of cognitive deficits in Ts mice, operated by selective pharmacology of α5-GABA_A_Rs (Braudeau et al., 2011; Duchon et al., 2019; Martínez-Cué et al., 2013). This subunit of GABA_A_Rs, whose synaptic vs. extrasynaptic expression is still debated (Ali and Thomson, 2008; Botta et al., 2015; Glykys and Mody, 2010; Hannan et al., 2020; Hausrat et al., 2015; Schulz et al., 2018; Serwanski et al., 2006), mediate dendritic synaptic inhibition from neocortical MCs (Ali and Thomson, 2008) and their hippocampal counterparts (the oriens-lacunoso moleculare or O-LM interneurons, Schulz et al., 2018). Here, we confirmed that α5-containg GABA_A_Rs are majorly responsible for MC-PN dendritic synaptic inhibition, due to the overall block of α5-IA, which was close to the max potency of this drug (Dawson et al., 2006).

The enhancement of the dendritic inhibitory loop involving MCs and PNs in Ts mice could not be attributed to alterations of axonal and/or dendritic arborizations in the mutant mice. On the other hand, it could be due to a combination of pre- and postsynaptic mechanisms, including alterations of release probability, number of release site or quantal size at either GABAergic or glutamatergic synapses involved in this circuit. Potentiation of glutamatergic synapses in Ts mice seem to be specific for PN-MC connections, as PN-PV synapses were similar in both genotypes, suggesting that presynaptic terminals of local glutamatergic synapses can undergo target-specific modulation of their strength. Intriguingly, it has been recently shown that an increase of the excitatory drive of hippocampal interneurons (due to triplication of GluR5 kainate receptor expression) could explain excess of inhibition received by pyramidal neurons in the Ts2Cje Down syndrome mouse model (Valbuena et al., 2019). Therefore, a similar gene overdose of kainate receptors can boost the recruitment of specific interneurons in the PFC of Ts mice.

The strong increase of membrane resistance of PV cells in Ts mice underlies the augmented intrinsic excitability and can produce early ectopic bouts of activity, thus contributing to network over-inhibition. This alteration of membrane resistance could be due to alterations in the expression of TWIK1 and TASK1 leak channels. Indeed, these channels underlie the developmental decrease of membrane resistance in PV cells (Okaty et al., 2009). In contrast, the inability of PV cells of Ts mice to sustain high frequency firing could prevent these interneurons from generating high-frequency bursts of action potentials and therefore have a detrimental effect on the temporal coding of these interneurons. Interestingly, however, despite the dramatic alterations of intrinsic excitability in Ts PV cells, their output synaptic perisomatic inhibition was similar in both genotypes. This, despite the widening of single PV-cell action potentials, which could, in principle change the presynaptic Ca^2+^ dynamics and thus alter release probability. Since action potentials are recorded in the soma, the lack of effect at PV-cell synapses could be due soma-specific alterations. Alternatively, since neocortical PV-PN GABAergic synapses exhibit high release probability (Deleuze et al., 2019; Kawaguchi and Kubota, 1998), alterations of spike width might not be enough to produce additional increases. Increase of action potential width in Ts mice could be due to changes in the expression of a different type of potassium channel, the Kv 3.1b. This potassium channel underlies the fast repolarization of action potentials and fast-spiking behavior of PV cells (Erisir et al., 1999). Future experiments will be necessary to pinpoint the exact molecular mechanism underlying the altered firing properties of PV cells, which, in Ts mice, have lost their characteristic fast-spiking signature.

Both potentiation of the dendritic inhibitory loop and increased PV-cell excitability are consistent with the alterations of PN spiking activity that we recorded *in vivo*. Indeed, reduced spike rates and increased tuning in the β-γ-frequency range are both consistent with increased activity of inhibitory neurons (Cardin et al., 2009; Sohal et al., 2009; Veit et al., 2017). The overall LFP power spectra observed in mPFC is similar in both Ts and Eu mice, consistently with a recent report (Chang et al., 2020). However, close examination of stLFP revealed a strong increase in the power of β- and low γ-frequency bands during periods of spiking activity. This likely reflects the consequences of alterations at the level of local microcircuits.

Importantly, it has been recently shown in the mouse visual cortex that in addition to the known role of PV cells in driving fast network oscillation, rhythmic activity in the β- and low γ-frequency bands also depends on SST interneurons (Veit et al., 2017). It is therefore possible that the specific alterations of MCs and PV interneurons shown here converge in sustaining the aberrant spiking and network activity observed in Ts mice.

In sum, here we report direct evidence for over-inhibition of mPFC circuits in a mouse model of DS. However, over-inhibition was not due to a generic increase of GABAergic signaling, but emerged from highly specific synaptic and intrinsic alterations of dendritic and somatic inhibitory circuits, respectively. Future experiments are necessary to reveal whether other inhibitory neuron types are also affected in DS. Likewise, it will be fundamental to assess whether specific dysfunctions of individual GABAergic circuits underlie different aspects of cognitive deficits (e.g. impaired memory and flexibility, autistic traits), which affect individuals with DS.

## Acknowledgements

We thank Vikaas Sohal, Nelson Rebola and Maria del Mar Dierssen Sotos for critically reading this manuscript. We also thank the ICM technical facilities PHENO-ICMICE and iGENSEQ.

This work was supported by “Investissements d’avenir” ANR-10-IAIHU-06, BBT-MOCONET; Agence Nationale de la Recherche (ANR-12-EMMA-0010; ANR-13-BSV4-0015-01; ANR-16-CE16-0007-02; ANR-17-CE16-0026-01; ANR-18-CE16-0001-01), Fondation Recherche Médicale (Equipe FRM DEQ20150331684 and EQU201903007860), NARSAD independent investigator grant and Fondation Lejeune (#1790).

## Author contributions

J.Z.S.M., M.C.P and A.B designed the experiments; J.Z.S.M, A.A., J.P. and C.D performed the experiments; J.Z.S.M., C.D., A.A., V.C. and J.P. analyzed the data; A.M.D.S and J.L. helped with *in vivo* recordings; M.C.P and A.B. supervised the project. All authors participated in writing the manuscript.

---

**Figure 1, figure supplement 1:**
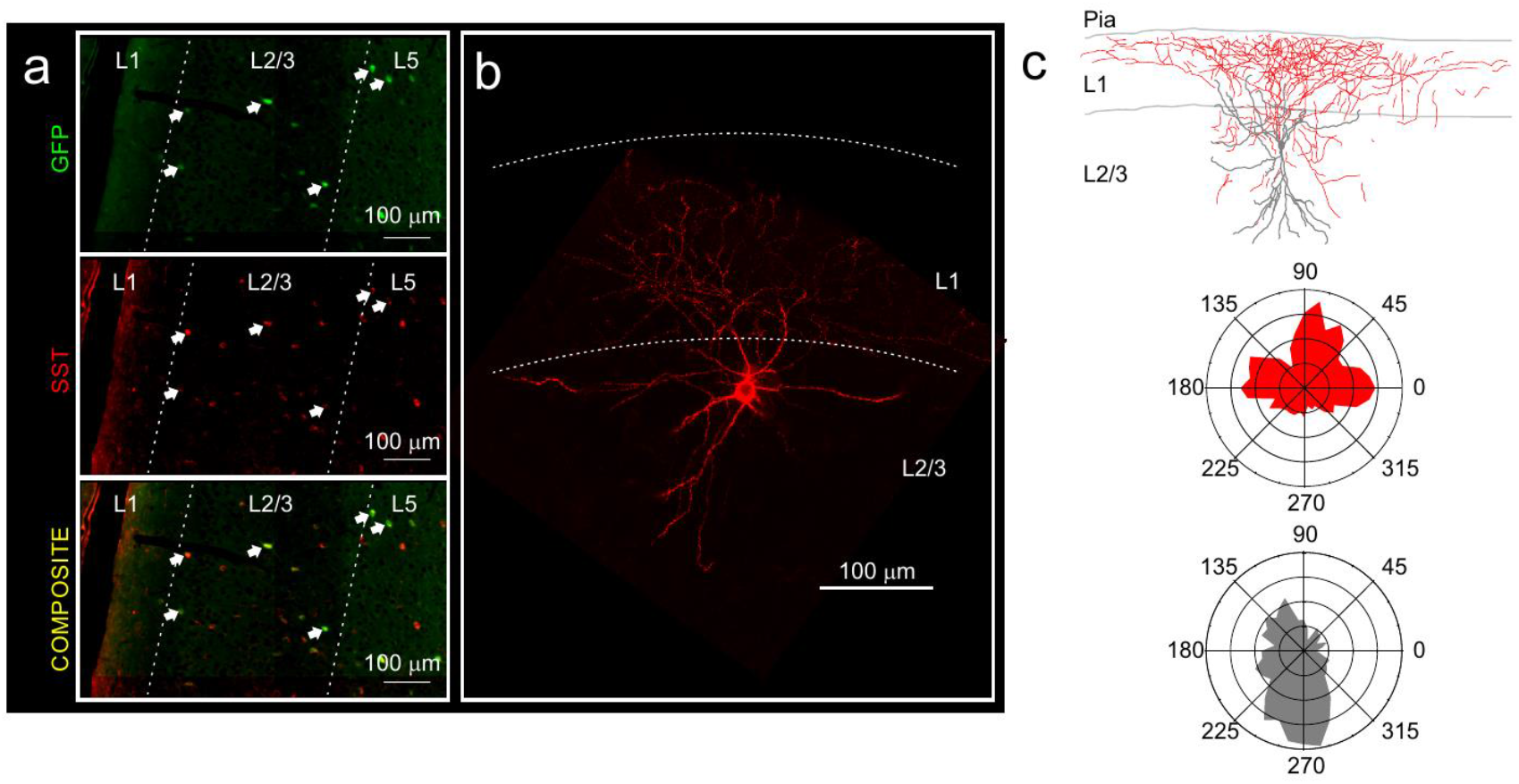
GFP positive interneurons in the prefrontal cortex of Ts::X98 mice are Martinotti cells. **a,** Immunostaining using anti-GFP (green) and anti-SST (red) antibodies in the mPFC. Note the co-localization of both markers. Arrows point to GFP and SST. **b,** Biocytin-filled GFP-positive interneuron revealed with Texas red coupled to streptavidin. Note the wide extension of the axonal plexus of Ts::X98 GFP-positive neurons in layer 1, typical of Martinotti cells. **c,** Top: Reconstructed morphology of the biocytin-filled cell of b and bottom: the polar histograms of its axonal (blue) and dendritic (pink) densities.

**Figure 2, figure supplement 1:**
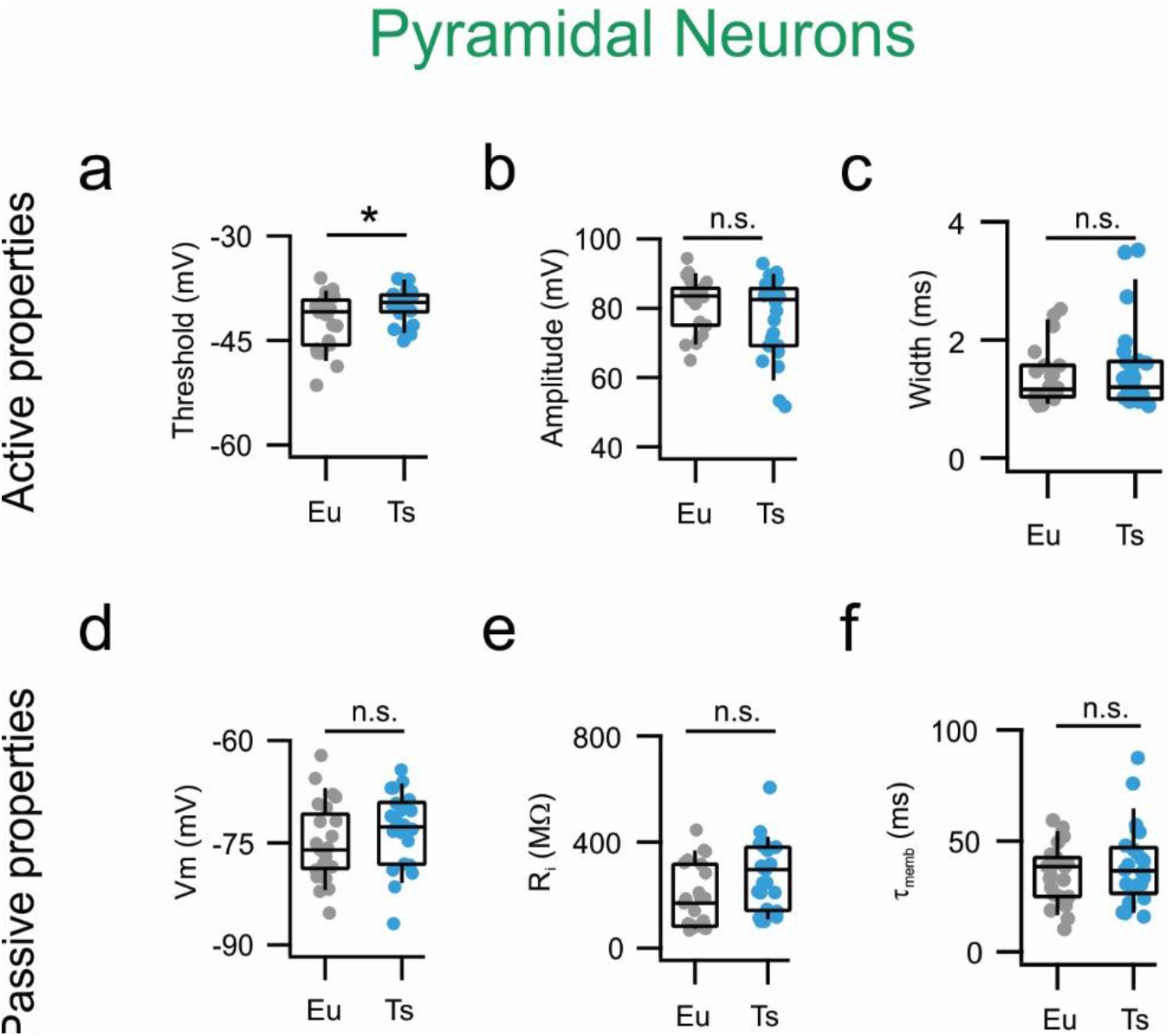
Single action potential and passive properties are normal Ts mPFC PNs. **a,** AP thresholds estimated as the voltage at which the derivative of Vm as function of time (dV/dt) reaches the limit of 10 mV/ms. Eu: −41 mV, −45-39 mV (n=23 cells) and Ts: −40, −41-39 mV mV (n=25 cells; Mann-Whitney U-test, p = 0.0453). **b,** AP amplitude measured as the difference between Vm at threshold and Vm at peak Eu: 84, 75-86 mV (n=23 cells) and Ts: 82, 69-85 mV (n=25 cells; Mann-Whitney U-test, p = 0.37486). **c,** AP width. Eu: 1.2, 1-1.6 ms (n=23 cells) and Ts: 1.2, 1-1.6 ms (n=25 cells; p = 0.74125, Mann-Whitney U-test). **d,** Resting membrane potential was also not significantly different between genotypes. Eu: −76, −79-72 mV (n=23 cells) and Ts: −73, −78-69 mV (n=25 cells; p = 0.74125, Mann-Whitney U-test). **e,** Input resistance, Eu: 255, 139-378 MΩ (n=23 cells), Ts: 178, 86-316 MΩ (n=25 cells, p=0.05918, Mann-Whitney U-test). **f,** Decay tau of the membrane Eu: 37, 28-48 (n=23 cells), Ts: 40, 27-47 (n=25 cells, p=0.90665, Mann-Whitney U-test). Boxplots: represent median, percentiles 25 and 75 and whiskers are percentiles 5 and 95. Individual points represent average values from individual cells.

**Figure 2, figure supplement 2:**
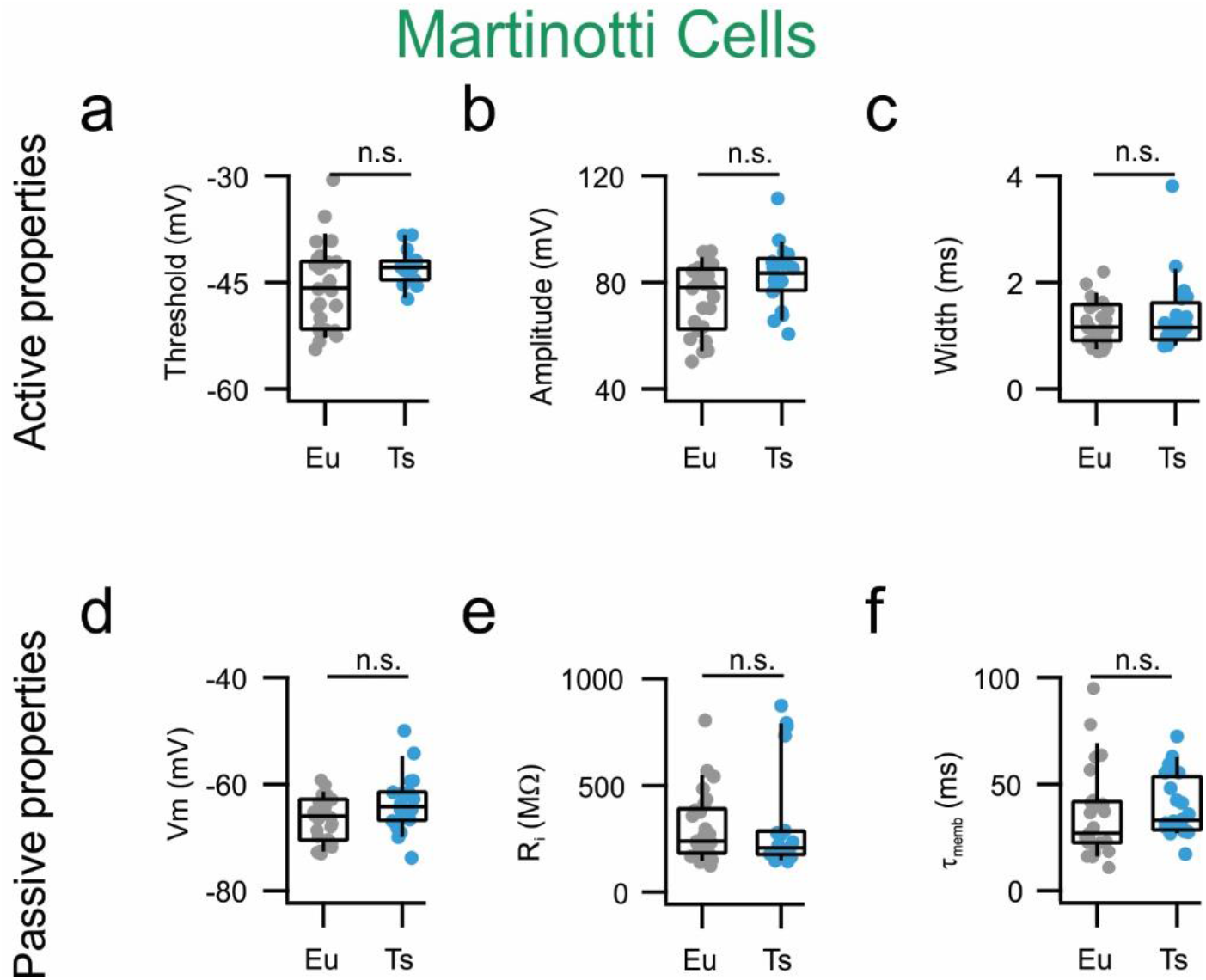
Active and passive properties are normal in mPFC MCs from Ts65Dn mice. **a,** AP thresholds estimated as the voltage at which the derivative of Vm as function of time (dV/dt) reaches the limit of 10 mV/ms. Eu: −46, −52-42 (n=23 cells); Ts: −43, −45-42 (n=25 cells, p=0.09877, Mann-Whitney U-test). **b,** AP amplitude measured as the difference between Vm at threshold and Vm at peak Eu: 79, 63-85 mV (n=26 cells), Ts: 82, 69-85 mV (n=20, p=0.07267, Mann-Whitney U-test). **c,** AP width. Eu: 1.2, 0.9-1.6 ms (n=26 cells), Ts: 1.2, 0.9-1.6 ms (n=20 cells, p=0.8680, Mann-Whitney U-test). **d,** Resting membrane potential. Eu: −66, −70-63 (n=26 cells), Ts: −64, −67-62 (n=20 cells, p=0.0566, Mann-Whitney U-test). **e,** Input resistance, Eu: 238, 183-382 MΩ (n=26 cells), Ts: 207, 176-282 MΩ (n=20 cells, p=0.57205, Mann-Whitney U-test). **f,** Decay tau of the membrane Eu: 27, 22-41 (n=26 cells), Ts: 33, 29-52 (n=20 cells, p=0.14067, Mann-Whitney U-test). Boxplots: represent median, percentiles 25 and 75 and whiskers are percentiles 5 and 95. Individual points represent average values from each recorded synaptic connection.

**Figure 2, figure supplement 3:**
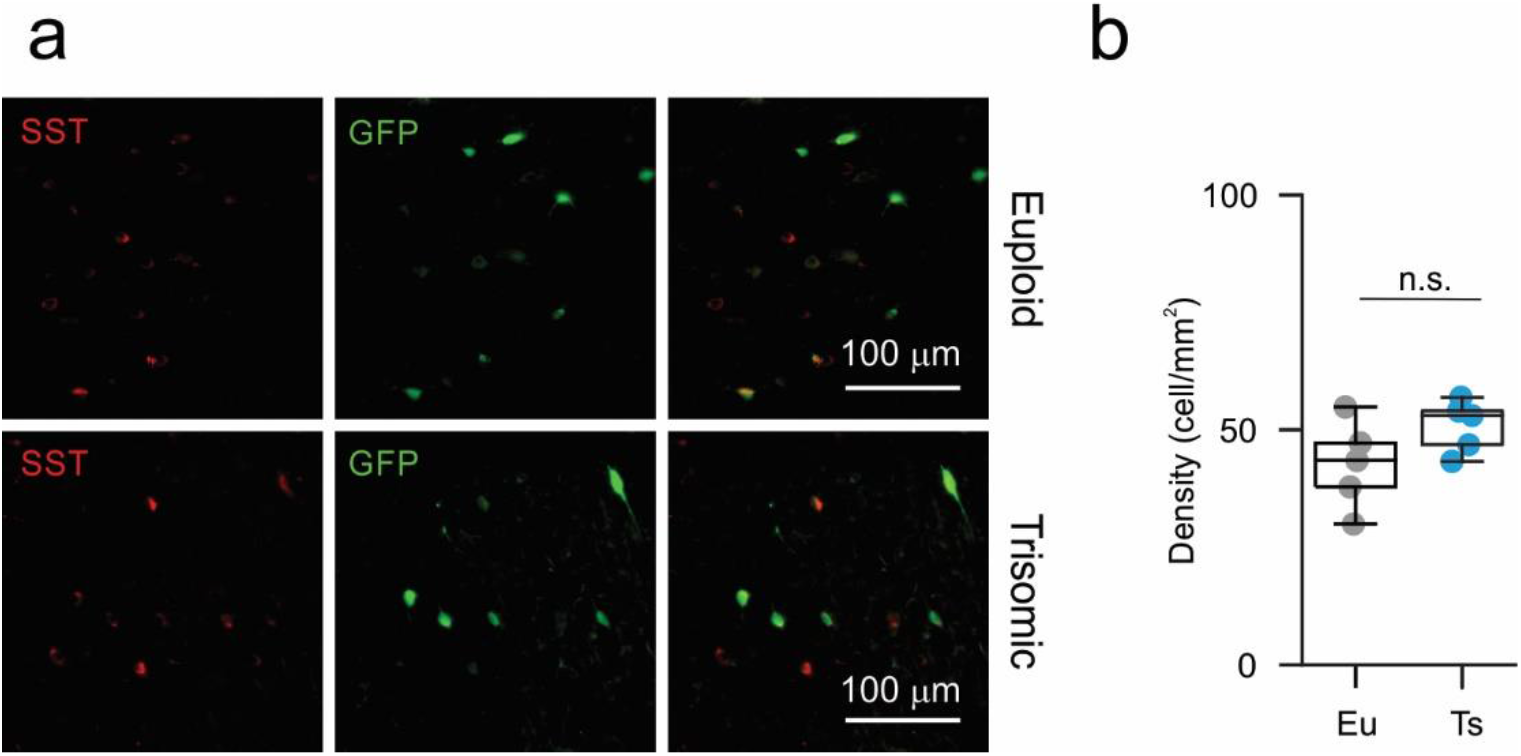
Normal SST-positive interneurons density in the mPFC of Ts mice. **a,** Representative fluorescence micrographs illustrating SST (red, left) and GFP (green, middle) immunostaining in Eu and Ts mice. Right column: merge of red and green channel b, Quantification of SST-positive cells density in mPFC L2/3 from Eu and Ts Ts::X98 mice.

**Figure 4, figure supplement 1:**
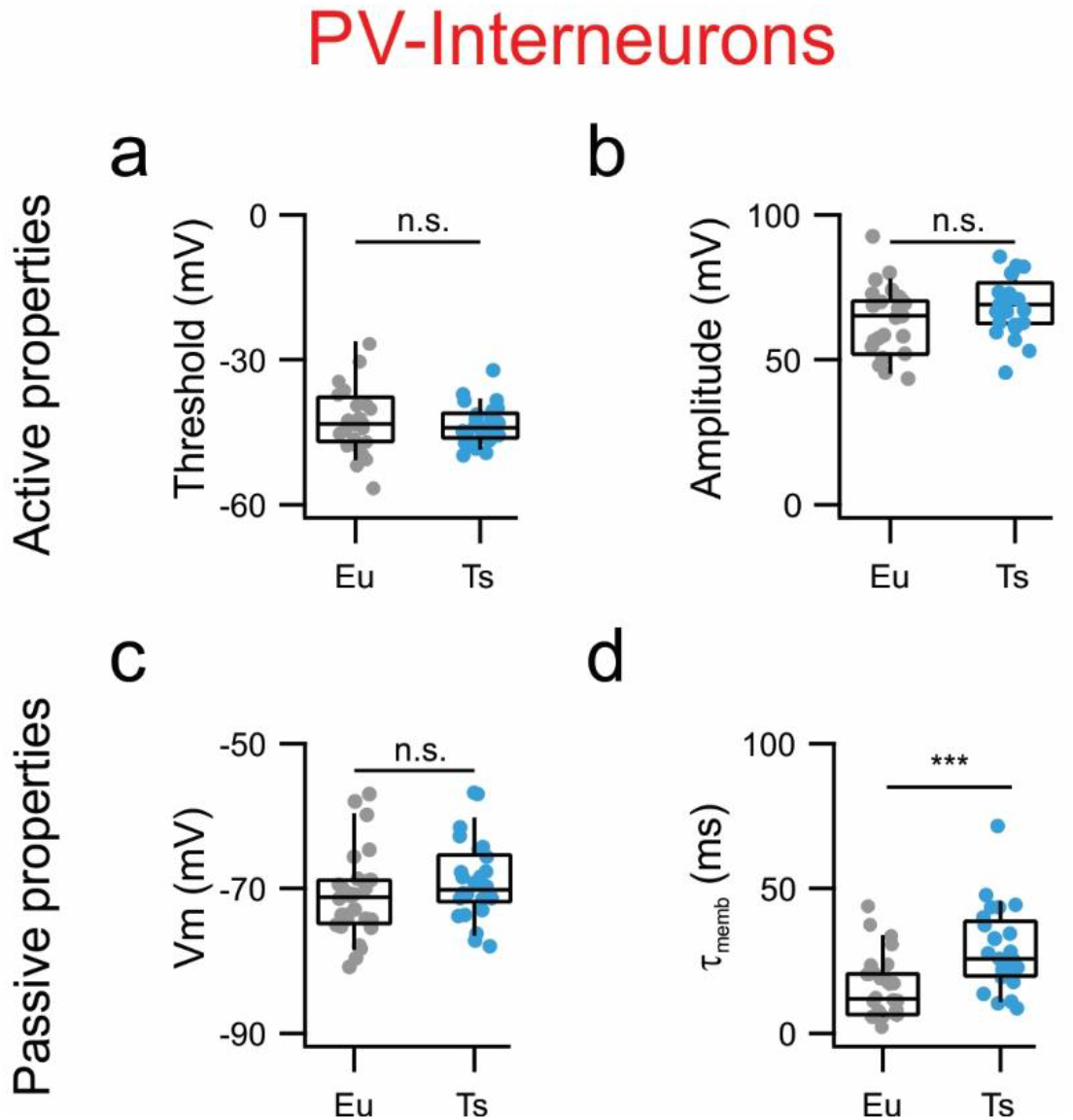
Specific alteration of mPFC PV-INs electrical properties in Ts65Dn mice. **a,** AP thresholds estimated as the voltage at which the derivative of Vm as function of time (dV/dt) reaches the limit of 10 mV/ms. Eu: −43 mV −47--39 (n=26 cells), Ts: −44 mV −46--41 (n=26 cells; p=0.0988 Student’s T test). **b,** AP amplitude measured as the difference between Vm at threshold and Vm at peak Eu: 79, 63-85 mV (n=26 cells), Ts: 69, 63-73 mV (n=26, p=0.8680, Mann-Whitney U-test). **c,** Resting membrane potential. Eu: −71, −75--69 (n=26 cells), Ts: −69, −71--66 (n=26 cells, p=0.0566, Student’s T test). **d,** Decay tau of the membrane Eu: 12, 7-21 ms (n=26 cells), Ts: 26, 20-37 ms (n=25 cells, p=0.14067, Mann-Whitney U-test). Boxplots: represent median, percentiles 25 and 75 and whiskers are percentiles 5 and 95. Individual points represent average values from each recorded synaptic connection.

**Figure 5, figure supplement 1:**
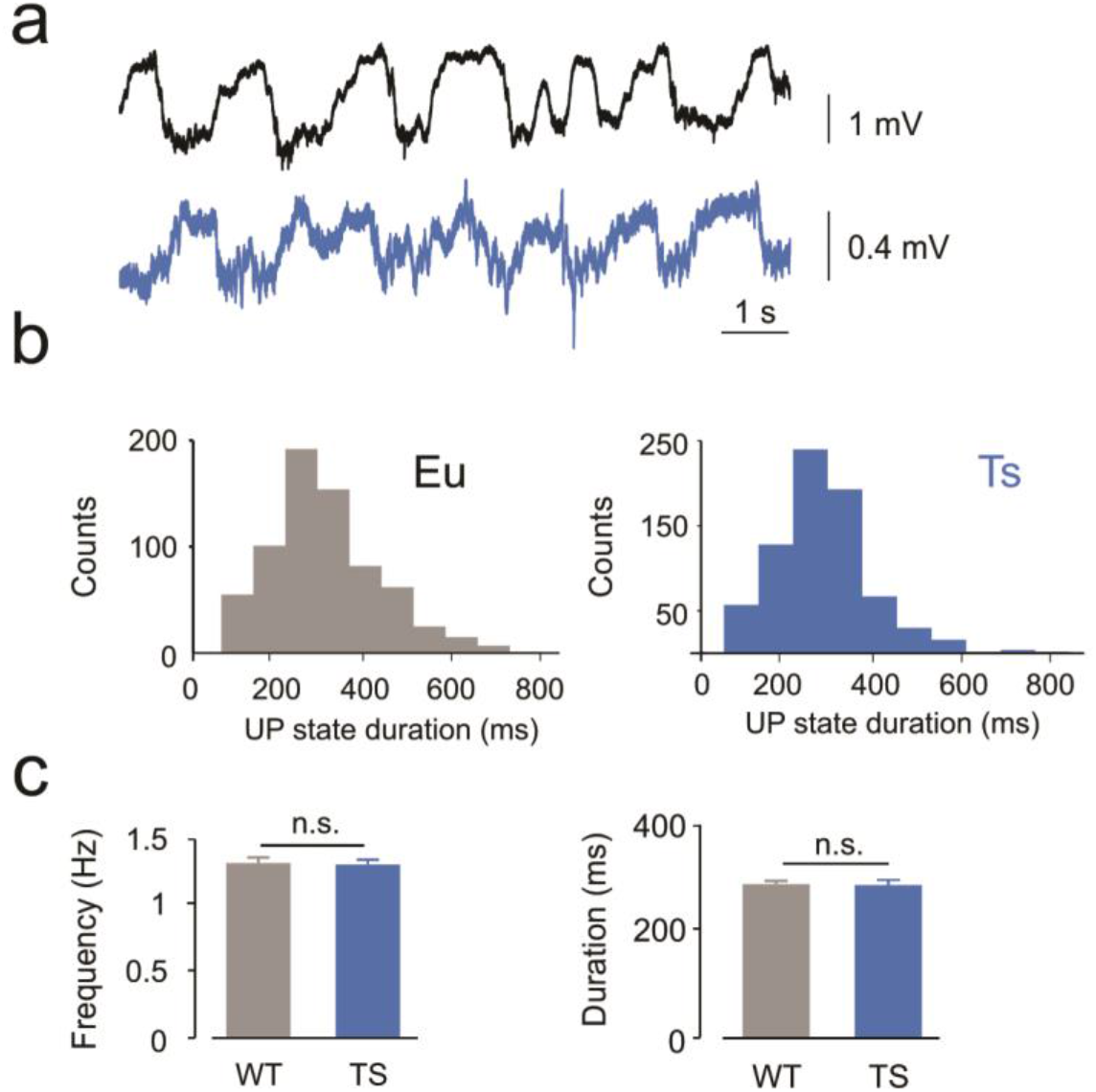
Characterization of slow wave oscillatory activity of LFP in anesthetized Eu and Ts mice. **a,** Representative LFP traces of anesthetized Eu (black, top) and Ts (bottom, blue) mice. **b,** Histograms of UP states duration from the examples shown in **a**, Eu (gray bars) and Ts (blue bars) examples. **c**, Average frequency, population data (left panel; Eu: 1.32 ± 0.02 Hz, Ts: 1.33 ± 0.02 Hz; Student’s T test, p= 0.86952) and average duration (right panel; Eu: 305.0 ± 3.3 ms, Ts: 310.9 ± 4.1 ms; Student’s T test, p=0.6352) of UP state did not significantly differed between genotypes (n=6 and 7 mice correspondingly).

## Methods

Experimental procedures followed national and European (2010/63/EU) guidelines, and have been approved by the authors’ institutional review boards and national authorities. All efforts were made to minimize suffering and reduce the number of animals. Experiments were performed on >4 week old and 18- to 25-day-old mice for *in vivo* and *ex vivo* recordings, respectively. We used *B6EiC3Sn a/A-Ts(17^16^)65Dn/J* (also known as Ts65Dn) mice (The Jackson Laboratory, Bar Harbor, Maine; stock #: 001924). For *ex vivo* slice experiments to label SST-positive MCs, Ts65Dn mice were crossed with *Tg(Gad1-EGFP)98Agmo/J*, also known as GAD67-GFP X98 or GFP-X98 (Jackson Laboratory). To label PV-INs, Ts65Dn mice were crossed with *C57BL/6-Tg(Pvalb-tdTomato)15Gfng/J (Pvalb-tdTomato*, Jackson Laboratory). Mice used in this study were of both sexes.

### In vivo LFP and juxtacellular recordings

Ts or Eu mice were anesthetized with 15% urethane (1.8 g/kg in physiological solution) and placed on a stereotaxic apparatus. The body temperature was constantly monitored and kept at 37 °C with a heating blanket. To ensure a deep and constant level of anesthesia, vibrissae movement, eyelid reflex, response to tail, and toe pinching were visually controlled before and during the surgery. A local lidocaine injection was performed over the cranial area of interest and, after a few minutes, a longitudinal incision was performed to expose the skull. Two small cranial windows (around 1 mm diameter) were opened at at 2.5 mm from bregma and ± 1 mm lateral to sagittal sinus (corresponding to the frontal lobe) carefully avoiding any damage to the main vessels while keeping the surface of the brain moist with the normal HEPES-buffered artificial cerebrospinal fluid. The dura was carefully removed using a metal needle. Glass pipettes used to record LFP had 1-2 MΩ resistance while those used for juxtacellular patch-clamp recordings typically had 5-7MΩ resistance. LFP and patch electrodes were pulled from borosilicate glass capillaries. Signals were amplified with a Multiclamp 700B patch-clamp amplifier (Molecular Devices), sampled at 20 KHz and filtered online at 10 KHz. Signals were digitized with a Digidata 1440A and acquired, using the pClamp 10 software package (Molecular Devices).

### Preparation of acute slices for electrophysiology

In order to record intrinsic and synaptic properties of L2/3 neurons of mPFC, we prepared acute cortical slices from the described mouse lines. For these experiments, we used slices cut in the coronal plane (300-350 μm thick). Animals were deeply anesthetized with saturating isofluorane (Vetflurane, Virbac) and immediately decapitated. The brain was then quickly removed (for groups of mice aged <P20, this procedure started right after deep anesthesia) and immersed in the cutting choline-based solution, containing the following (in mM): 126 choline chloride, 16 glucose, 26 NaHCO_3_, 2.5 KCl, 1.25 NaH_2_PO_4_, 7 MgSO_4_, 0.5 CaCl_2_, cooled to 4°C and equilibrated with a 95-5% O_2_-CO_2_ gas mixture. Slices were cut with a vibratome (Leica VT1200S) in cutting solution and then incubated in oxygenated artificial cerebrospinal fluid (aCSF) composed of (in mM): 126 NaCl, 20 glucose, 26 NaHCO_3_, 2.5 KCl, 1.25 NaH_2_PO_4_, 1 MgSO_4_, 2 CaCl_2_ (pH 7.35, 310-320mOsm/L), initially at 34°C for 30 min, and subsequently at room temperature, before being transferred to the recording chamber where recordings were obtained at 30–32°C.

### Slice electrophysiology

Whole-cell patch-clamp recordings were performed in L2/3 of the medial prefrontal cortex (mPFC) neurons. Inhibitory PV-expressing interneurons, labeled with TdTomato in Ts65Dn mice crossed with *Pvalb-tdTomato* mice and Martinotti cells, labeled with GFP in Ts65Dn crossed with GFP-X98 mice, were identified using LED illumination (OptoLED, Cairn Research, Faversham, UK). Excitatory pyramidal neurons (PNs) were visually identified using infrared video microscopy, as cells lacking expression of fluorescent proteins and with somatas exhibiting the classical pyramidal shape. Accordingly, when depolarized with DC current pulses PNs exhibited a typical firing pattern of regular-spiking cells. We used different intracellular solutions depending on the type of experiment and the nature of the responses we wanted to assess. To study passive properties, intrinsic excitability, AP waveform and glutamatergic spontaneous transmission, electrodes were filled with an intracellular solution containing (in mM): 127 K-gluconate, 6 KCl, 10 Hepes, 1 EGTA, 2 MgCl2, 4 Mg-ATP, 0.3 Na-GTP; pH adjusted to 7.3 with KOH; 290–300 mOsm. The estimated reversal potential for chloride (E_Cl_) was approximately −69 mV based on the Nernst equation. To measure GABAergic currents elicited by perisomatic-targeting interneurons, PNs were patched using an intracellular solution containing (in mM): 65 K-gluconate, 70 KCl, 10 Hepes, 1 EGTA, 2 MgCl2, 4 Mg-ATP, 0.3 Na-GTP; pH adjusted to 7.3 with KOH; 290-300 mOsm (the estimated ECl was approximately −16 mV based on the Nernst equation). For distal dendritic uIPSCs, we used a cesium-based solution containing (in mM): 145 CsCl, 10 Hepes, 1 EGTA, 0.1 CaCl2, 2 MgCl2, 4.6 Mg-ATP, 0.4 Na-GTP, 5 QX314-Cl; pH adjusted to 7.3 with CsOH; 290–300 mOsm. Under these recording conditions, activation of GABA_A_ receptors resulted in inward currents at a holding potential (Vh) of −70 mV. Voltage values were not corrected for liquid junction potential. Patch electrodes were pulled from borosilicate glass capillaries and had a typical tip resistance of 2–3 MΩ. Signals were amplified with a Multiclamp 700B patch-clamp amplifier (Molecular Devices), sampled at 20–50 KHz and filtered at 4 KHz (for voltage-clamp experiments) and 10 KHz (for current-clamp experiments). Signals were digitized with a Digidata 1440A and acquired, using the pClamp 10 software package (Molecular Devices).

*For paired recordings*, unitary synaptic responses were elicited in voltage-clamp mode by brief somatic depolarizing steps (−70 to 0 mV, 1-2 ms) evoking action currents in presynaptic cells. Neurons were held at −70 mV and a train of 5 presynaptic spikes at 50 Hz was applied.

### Immunohistochemistry

Parvalbumin, SST and GFP staining were performed on 20-50 μm-thick slices. Briefly, mice were perfused with 0.9% NaCl solution containing Heparin and 4% paraformaldehyde (PFA). Brains were cryo-protected by placing them overnight in 30% sucrose solution and then frozen in Isopentane at a temperature <−50°C. Brains were sliced with a freezing microtome (ThermoFisher HM450). Permeabilization in a blocking solution of PBT with 0.3% Triton and 10% Normal Goat Serum was done at room temperature for 2 hr. Slices were then incubated overnight (4°C) in the same blocking solution containing the primary rabbit anti-PV antibody (1:1000; Thermo Scientific) and mouse anti-SST antibody (1:250; Santa Cruz Biotechnologies). Slices were then rinsed three times in PBS (10 min each) at room temperature and incubated with goat anti-rabbit and a goat anti-mouse antibody (1:500; Jackson IR) coupled to Alexa-488 or 633 for 3.5 hr at room temperature. Slices were then rinsed three times in PBS (10 min each) at room temperature and coverslipped in mounting medium (Fluoromount, Sigma Aldrich F4680). Immunofluorescence was then observed with a slide scanner (Zeiss, Axio Scan.Z1).

### Morphological reconstruction

Biocytin Fills: To reliably reconstruct the fine axonal branches of cortical neurons, dedicated experiments were performed following the classical avidin-biotin-peroxidase method. Biocytin (Sigma) was added to the intracellular solution at a high concentration (5-10mg/ml), which required extensive sonication. At the end of recordings, the patch pipette was removed carefully until obtaining an inside out patch. The slice was then left in the recording chamber for at least further 5-10 min to allow further diffusion. Slices were then fixed with 4% paraformaldehyde in phosphate buffer saline (PBS, Sigma) for at least 48 h. Following fixation, slices were incubated with the avidin-biotin complex (Vector Labs) and a high concentration of detergent (Triton-X100, 5%) for at least two days before staining with 3,3’Diaminobenzidine (DAB, AbCam). Cells were then reconstructed and cortical layers delimited using Neurolucida 7 (MBF Bioscience) and the most up to date mouse atlas (Allen Institute).

### Data analysis

Electrophysiological and statistical analysis was performed using built-in and custom-written routines made for Igor Pro (WaveMetrics, Lake Oswego, OR, USA), MATLAB R2017b 9.3.0.713579 Natick, Massachusetts: The MathWorks Inc.; Origin (Pro) 2016 OriginLab Corporation, Northampton, MA, USA; Prism version 7.00 for Windows, GraphPad Software, La Jolla California USA; and Python Software Foundation. Python Language Reference, version 3.6, available at http://www.python.org.

#### Analysis of in vivo recordings

Traces obtained from juxtacellular recording were high pass filtered (cutoff: 5 Hz) and spikes were detected based on threshold=1.5 mV. Spike rate was estimated as the total number of spikes detected divided by the total duration of the recording. The peak time (tpeak) corresponding to each detected action potential was used to select the segment of LFP between: [t_peak_-100 ms to tpeak+100 ms]. The instantaneous phase was estimated using Hilbert transform on decimated LFP and subsequently used to estimate phase locking. For LFP analysis, extracellular potentials were down-sampled (1 kHz) and low-pass filtered (cutoff frequency, 100 Hz). Power spectra were generated using a Hann window (window length: 4096 points, 50% overlap).

Phase locking was determined using pairwise phase consistency (PPC) estimation, defined as:

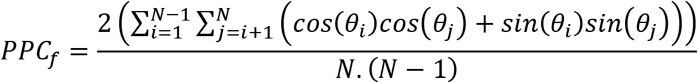

Where N is the total number of action potentials and θ_*i*_ is the phase of the *i*^th^ spike and θ_*j*_ the *j^th^*.

#### UP and DOWN states detection

Cortical states were detected as described elsewhere^27^. Briefly, filtered and decimated LFP was used to calculate UP and DOWN state likelihood decision (or evidence) variable (S_comb_) based on low frequency (<4Hz) oscillation phase and high frequencies (20–100Hz) composition of the LFP. To determine the thresholds to detect different cortical states (UP, Intermediate, DOWN) states, the distribution of the combined evidence variable,*S_comb_*, was fitted by a mixture of three Gaussians, each representing their corresponding cortical state, UP (highest level of the signal), Intermediate (Intermediate level) and DOWN (lowest level of the signal). Periods of the combined signal, *S_comb_*, that were above the UP threshold, *μ_UP-LFP_* − 3 * *σ_UP-LFP_*, were considered the periods of UP states. Similarly, the periods below the DOWN threshold, *μ_D0WN-LFP_* − 3 * *σ_D0WN-LFP_*, were considered the periods of DOWN states (means and variances of the Gaussians are represented as *μ_UP_, μ_DOWN_*, and *σ_UP_, σ_DOWN_* for the up and down cortical states, respectively). Periods of UP and DOWN states were refined further by putting constraints on the interval between two states and duration of a state. Minimum interval between two states and duration of a state were set 50 and 70 ms, respectively.

#### Analysis of slices electrophysiological recordings

Input resistance (R_i_) was estimated as the slope of the current to voltage relationship obtained with upon the injection of −25, 0 and 25 pA to a cell kept at resting potential. Membrane time constant was measured estimated fitting the time course of V_memb_ after the injection of a 2s, −25 pA current step.

We used protocols of increasing steps of current injection (−50 to 500 pA in steps of 25 or 50 pA and 2 seconds duration). Action potentials were detected using a threshold based routine. Threshold was set at 0 mV. Firing dynamics was evaluated fitting F-I curves from individual cells to a sigmoidal function:

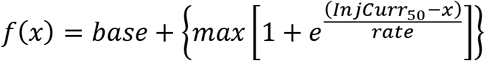

Where *base:* minimal spiking rate, *max:* maximal spiking rate achieved, InjCurr_50_: current injected necessary to reach 50% of *max*, and *rate*: the gain. The first action potential evoked was taken to measure amplitude, width and threshold. Threshold was considered as the potential at which dV/dt reached 10 mV/ms; amplitude was the difference between peak amplitude and threshold and half-width was the time interval between rise and decay phase measured at 50% of amplitude.

Action potential threshold was defined as the membrane potential value (V_m_) at which dV/dt become bigger than 10 mV/ms. Action potential amplitude is: *Amp = Amp_peak_* − *threshold*where Amp_peak_ is the AP peak potential. Action potential width was measured at 50% of amplitude.

### Statistical analysis

Normal distribution of samples was systematically assessed (Shapiro-Wilkinson normality test). Normal distributed samples were statistically compared using two-tailed Student’s *t* test unless otherwise stated. When data distribution was not normal we used two-tailed Mann Whitney U-test. Compiled data are reported and presented as whisker box plots (the upper and lower whiskers representing the 90th and 10th percentiles, respectively, and the upper and lower boxes representing the 75th and 25th percentiles, respectively, and the horizontal line representing the median or the mean ± s.e.m., with single data points plotted. Differences were considered significant if p<0.05 (*p<0.05, **p<0.01, ***p<0.001).

## Supplementary Tables

### Dendritic inhibition

Evaluation of synaptic efficiency in the dendritic inhibitory loop composed by MCs and PNs.

**Supplementary table 1:** MC to PN synaptic efficiency evaluation.

|  |  | Dendritic Inhibition |  |  |  |  |  |  |  |  |  |
| --- | --- | --- | --- | --- | --- | --- | --- | --- | --- | --- | --- |
|  |  | Euploid |  |  |  | Ts65Dn |  |  |  |  |  |
|  |  | median | Q1 | Q3 | n | median | Q1 | Q3 | n | test | p value |
| **MC-PN IPSC** | Amplitude (pA) | 29.89 | 20.24 | 52.98 | 11 | 97.09 | 54.49 | 135.86 | 15 | MW-test | 0.0095 |
|  | Failure Rate | 0.20 | 0.11 | 0.25 | 11 | 0.00 | 0.00 | 0.13 | 15 | MW-test | 0.0201 |
|  | Charge (pC) | 1.28 | 0.96 | 1.84 | 11 | 6.86 | 2.96 | 8.32 | 15 | MW-test | 0.00595 |
| | α5IA % block | 41 | 24.20 | 57.20 | 4 | 57.4 | 33.10 | 73.10 | 6 | MW-test | 0.2278 |

Median: quantile 50; Q1: quantile 25; Q3: quantile 75; n: number of cells; IPSC: Inhibitory postsynaptic current.

**Supplementary table 2:**
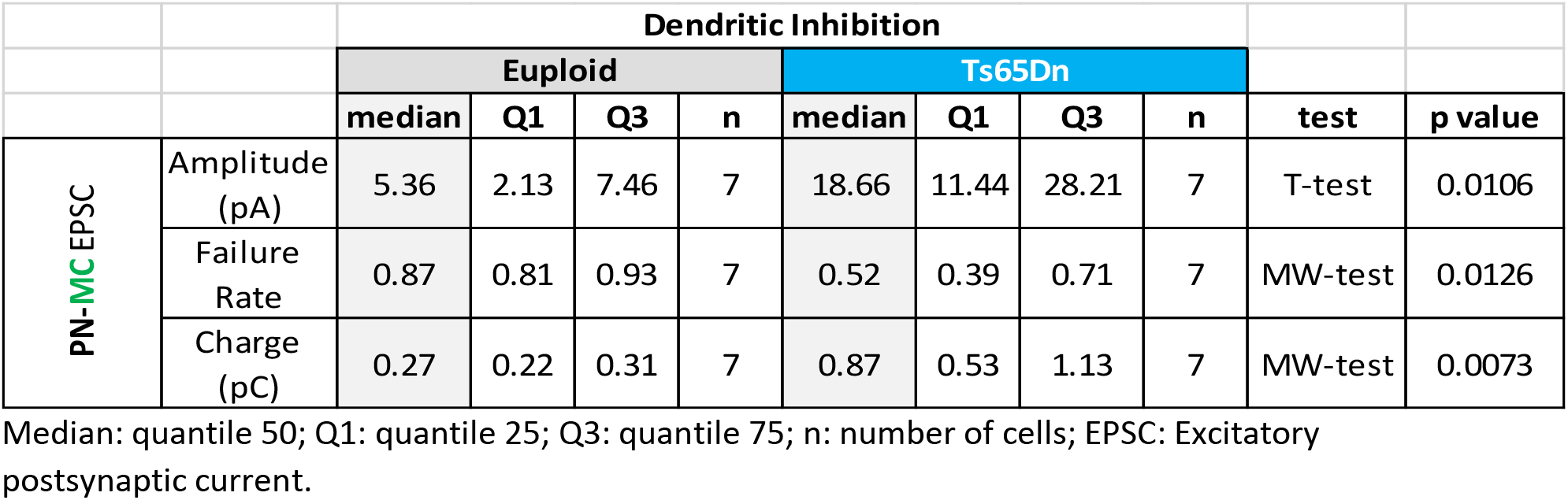
PN to MC synaptic efficiency evaluation.

**Supplementary table 3:**
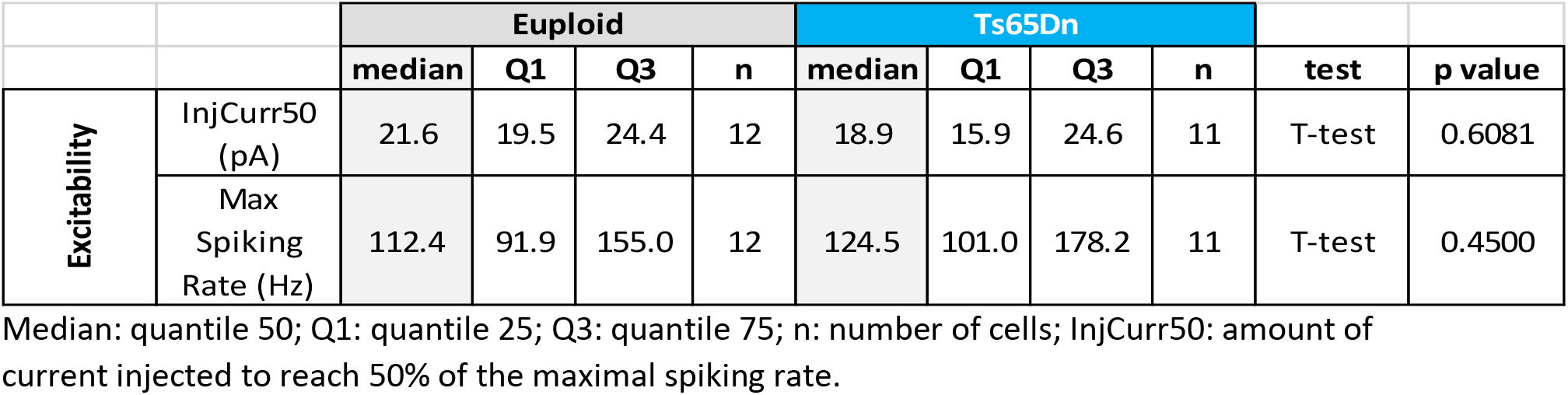
PNs excitability.

|  |  | Euploid |  |  |  | Ts65Dn |  |  |  | test | p value |
| --- | --- | --- | --- | --- | --- | --- | --- | --- | --- | --- | --- |
|  |  | median | Q1 | Q3 | n | median | Q1 | Q3 | n |  |  |
| Excitability | InjCurr50 (pA) | 21.6 | 19.5 | 24.4 | 12 | 18.9 | 15.9 | 24.6 | 11 | T-test | 0.6081 |
|  | Max Spiking Rate (Hz) | 112.4 | 91.9 | 155.0 | 12 | 124.5 | 101.0 | 178.2 | 11 | T-test | 0.4500 |

Median: quantile 50; Q1: quantile 25; Q3: quantile 75; n: number of cells; InjCurr50: amount of current injected to reach 50% of the maximal spiking rate.

**Supplementary table 4:** MCs excitability.

|  |  | Euploid |  |  |  | Ts65Dn |  |  |  | test | p value |
| --- | --- | --- | --- | --- | --- | --- | --- | --- | --- | --- | --- |
|  |  | median | Q1 | Q3 | n | median | Q1 | Q3 | n |  |  |
| Excitability | InjCurr50 (pA) | 40.9 | 26.9 | 50.4 | 25 | 48.9 | 25.4 | 70.2 | 20 | MW-test | 0.6073 |
|  | Max Spiking Rate (Hz) | 101.8 | 68.3 | 124.0 | 25 | 86.5 | 63.2 | 109.7 | 20 | T-test | 0.4436 |

Median: quantile 50; Q1: quantile 25; Q3: quantile 75; n: number of cells; InjCurr50: amount of current injected to reach 50% of the maximal spiking rate.

**Supplementary table 5:** PNs passive and action potential (AP) properties.

|  |  | Euploid |  |  |  | Ts65Dn |  |  |  | test | p value |
| --- | --- | --- | --- | --- | --- | --- | --- | --- | --- | --- | --- |
|  |  | median | Q1 | Q3 | n | median | Q1 | Q3 | n |  |  |
| Passive properties | Vrest (mV) | -76 | -79 | -72 | 25 | -73 | -78 | -69 | 22 | T-test | 0.2473 |
|  | Ri (MΩ) | 255 | 139 | 378 | 25 | 178 | 86 | 316 | 22 | T-test | 0.7185 |
|  | Tau memb (ms) | 36.5 | 28.1 | 47.9 | 25 | 39.7 | 26.8 | 46.9 | 22 | T-test | 0.5790 |
| AP properties | Threshold (mV) | -40.9 | -45.6 | -39.2 | 25 | -39.6 | -40.8 | -38.5 | 22 | MW-test | 0.0453 |
|  | Amplitude (mV) | 83.6 | 75.1 | 85.8 | 25 | 82.5 | 69.2 | 85.5 | 22 | MW-test | 0.3749 |
|  | Width (ms) | 1.2 | 1.0 | 1.6 | 25 | 1.2 | 1.0 | 1.6 | 22 | MW-test | 0.7413 |

Median: quantile 50; Q1: quantile 25; Q3: quantile 75; n: number of cells; Vrest: Resting membrane potential; Ri : input resistance.

**Supplementary table 6:** MCs passive and action potential (AP) properties.

|  |  | Euploid |  |  |  | Ts65Dn |  |  |  | test | p value |
| --- | --- | --- | --- | --- | --- | --- | --- | --- | --- | --- | --- |
|  |  | median | Q1 | Q3 | n | median | Q1 | Q3 | n |  |  |
| Passive properties | Vrest (mV) | -66 | -70 | -63 | 26 | -64 | -67 | -62 | 20 | T-test | 0.0566 |
|  | Ri (MΩ) | 238 | 183 | 382 | 26 | 207 | 176 | 282 | 20 | MW-test | 0.5721 |
|  | Tau memb (ms) | 27.0 | 22.4 | 41.4 | 25 | 33.0 | 28.7 | 51.7 | 20 | MW-test | 0.1407 |
| AP properties | Threshold (mV) | 79.0 | 63.2 | 85.1 | 26 | 83.4 | 77.3 | 88.5 | 20 | MW-test | 0.0988 |
|  | Amplitude (mV) | -45.8 | -51.5 | -42.2 | 26 | -42.9 | -44.6 | -42.0 | 20 | MW-test | 0.0727 |
|  | Width (ms) | 1.2 | 0.9 | 1.6 | 26 | 1.2 | 0.9 | 1.5 | 20 | MW-test | 0.8680 |

Median: quantile 50; Q1: quantile 25; Q3: quantile 75; n: number of cells; Vrest: Resting membrane potential ; Ri : input resistance.

### Perisomatic inhibition

Evaluation of synaptic efficiency in the perisomatic inhibitory loop composed by PVs and PNs.

**Supplementary table 7:** Direct perisomatic inhibition. PV to PN synaptic efficiency evaluation.

|  |  | Perisomatic inhibition |  |  |  |  |  |  |  |  |  |
| --- | --- | --- | --- | --- | --- | --- | --- | --- | --- | --- | --- |
|  |  | Euploid |  |  |  | Ts65Dn |  |  |  |  |  |
|  |  | median | Q1 | Q3 | n | median | Q1 | Q3 | n | test | p value |
| PV-PN IPSC | Amplitude (pA) | 188.43 | 74.87 | 289.03 | 26 | 144.52 | 46.10 | 209.78 | 15 | MW-test | 0.49 |
|  | Failure Rate | 0.00 | 0.00 | 0.03 | 26 | 0.00 | 0.00 | 0.00 | 15 | MW-test | 0.7185 |
|  | Charge (pC) | 2.88 | 1.92 | 5.02 | 26 | 4.89 | 1.89 | 5.85 | 15 | MW-test | 0.579 |

Median: quantile 50; Q1: quantile 25; Q3: quantile 75; n: number of cells; IPSC: Inhibitory postsynaptic current.

**Supplementary table 8:** Recruitment of PV-INs by PNs. PN to PV synaptic efficiency evaluation.

|  |  | Perisomatic inhibition |  |  |  |  |  |  |  |  |  |
| --- | --- | --- | --- | --- | --- | --- | --- | --- | --- | --- | --- |
|  |  | Euploid |  |  |  | Ts65Dn |  |  |  |  |  |
|  |  | median | Q1 | Q3 | n | median | Q1 | Q3 | n | test | p value |
| PN-PV EPSC | Amplitude (pA) | 65.66 | 19.28 | 124.50 | 15 | 20.41 | 14.10 | 84.25 | 9 | MW-test | 0.23304 |
|  | Failure Rate | 0.00 | 0.00 | 0.27 | 15 | 0.13 | 0.00 | 0.57 | 9 | MW-test | 0.21402 |
|  | Charge (pC) | 0.62 | 0.29 | 0.90 | 15 | 0.42 | 0.20 | 0.55 | 9 | MW-test | 0.3711 |

Median: quantile 50; Q1: quantile 25; Q3: quantile 75; n: number of cells; EPSC: Excitatory postsynaptic current.

**Supplementary table 9:** PV-INs passive and action potential (AP) properties.

|  |  | Euploid |  |  |  | Ts65Dn |  |  |  | test | p value |
| --- | --- | --- | --- | --- | --- | --- | --- | --- | --- | --- | --- |
|  |  | median | Q1 | Q3 | n | median | Q1 | Q3 | n |  |  |
| Excitability | InjCurr50 (pA) | 310.4 | 136.3 | 394.6 | 26 | 92.9 | 63.5 | 142.7 | 26 | MW-test | 0.0003 |
|  | Max Spiking Rate (Hz) | 99.5 | 45.2 | 142.3 | 26 | 47.9 | 45.3 | 63.3 | 26 | MW-test | 0.0001 |

Median: quantile 50; Q1: quantile 25; Q3: quantile 75; n: number of cells; InjCurr50: amount of current injected to reach 50% of the maximal spiking rate.

**Supplementary table 10:** PV-INs passive and action potential (AP) properties.

|  |  | Euploid |  |  |  | Ts65Dn |  |  |  | test | p value |
| --- | --- | --- | --- | --- | --- | --- | --- | --- | --- | --- | --- |
|  |  | median | Q1 | Q3 | n | median | Q1 | Q3 | n |  |  |
| Passive properties | Vrest (mV) | -71 | -75 | -69 | 28 | -70 | -71 | -66 | 26 | T-test | 0.1808 |
|  | Ri (MΩ) | 154 | 93 | 271 | 28 | 406 | 264 | 581 | 26 | MW-test | 0.0001 |
|  | Tau memb (ms) | 12.0 | 6.5 | 20.6 | 28 | 25.8 | 20.2 | 37.3 | 25 | MW-test | 0.0003 |
| AP properties | Threshold (mV) | -43.4 | -47.0 | -39.5 | 26 | -44.0 | -46.0 | -41.3 | 26 | T-test | 0.7891 |
|  | Amplitude (mV) | 65.2 | 54.6 | 70.3 | 25 | 69.0 | 62.8 | 73.4 | 25 | T-test | 0.1019 |
|  | Width (ms) | 0.8 | 0.7 | 1.3 | 26 | 1.3 | 1.0 | 1.6 | 26 | MW-test | 0.0012 |

Median: quantile 50; Q1: quantile 25; Q3: quantile 75; n: number of cells; Vrest: Resting membrane potential; Ri : input resistance.

### Ts65Dn in vivo activity

**Supplementary table 11:** LFP and single cell spiking recorded in vivo.

|  |  | Euploid |  |  |  |  | Ts65Dn |  |  |  |  |  |  |
| --- | --- | --- | --- | --- | --- | --- | --- | --- | --- | --- | --- | --- | --- |
|  |  | median | Q1 | Q3 | N cells | N mice | median | Q1 | Q3 | N cells | N mice | test | p value |
|  | Spiking Rate (Hz) | 0.81 | 0.56 | 1.24 | 28 | 6 | 0.42 | 0.22 | 0.92 | 23 | 5 | MW-test | 0.0348 |
| stLFP | peak-to-peak Amp (μV) | 7.3 | 6.0 | 8.6 | 22 | 6 | 13.3 | 12.3 | 17.1 | 11 | 5 | MW-test | 0.0011 |

Median: quantile 50; Q1: quantile 25; Q3: quantile 75; n: number of cells.

**Supplementary table 12:** Pairwise Phase Consistency (PPC) descriptive statistics and hypothesis tests between genotypes for each frequency band analyzed.

|  |  | Euploid |  |  |  |  | Ts65Dn |  |  |  |  |  |  |
| --- | --- | --- | --- | --- | --- | --- | --- | --- | --- | --- | --- | --- | --- |
|  | Freq. band | median | Q1 | Q3 | N cells | N mice | median | Q1 | Q3 | N cells | N mice | test | p value |
| PPC | 4-10 Hz | 0.0085 | 0.0053 | 0.0242 | 22 | 6 | 0.0106 | 0.0064 | 0.0164 | 11 | 5 | MW-test | 0.4270 |
|  | 10-20 Hz | 0.0046 | 0.0011 | 0.0237 | 22 | 6 | 0.0213 | 0.0163 | 0.0379 | 11 | 5 | MW-test | 0.0215 |
|  | 20-40 Hz | 0.0024 | -0.0002 | 0.0085 | 22 | 6 | 0.0066 | 0.0059 | 0.0193 | 11 | 5 | MW-test | 0.0418 |
|  | 40-60 Hz | 0.0009 | -0.0009 | 0.0025 | 22 | 6 | 0.0052 | 0.0019 | 0.0079 | 11 | 5 | MW-test | 0.0164 |
|  | 60-80 Hz | -0.0003 | -0.0007 | 0.0004 | 22 | 6 | 0.0001 | -0.0005 | 0.0009 | 11 | 5 | MW-test | 0.2308 |
|  | 80-100 Hz | -0.0002 | -0.0009 | 0.0013 | 22 | 6 | -0.0004 | -0.0008 | 0.0002 | 11 | 5 | MW-test | 0.3161 |

Median: quantile 50; Q1: quantile 25; Q3: quantile 75; n: number of cells. We considered only cells from which we recorded at least 250 action potentials in order to obtain reliable PPC values.

## Notes

### Competing Interest Statement

The authors have declared no competing interest.

## References

Abbas AI, Sundiang MJM, Henoch B, Morton MP, Bolkan SS, Park AJ, Harris AZ, Kellendonk C, Gordon JA. 2018. Somatostatin Interneurons Facilitate Hippocampal-Prefrontal Synchrony and Prefrontal Spatial Encoding. Neuron 100:926–939.e3. doi:10.1016/j.neuron.2018.09.029

Ali AB, Thomson AM. 2008. Synaptic a5 subunit-containing GABA_A_ receptors mediate ipsps elicited by dendrite-preferring cells in rat neocortex. Cereb Cortex 18:1260–1271. doi:10.1093/cercor/bhm160

Antonarakis SE, Skotko BG, Rafii MS, Strydom A, Pape SE, Bianchi DW, Sherman SL, Reeves RH. 2020. Down syndrome. Nat Rev Dis Prim 6. doi:10.1038/s41572-019-0143-7

Botta P, Demmou L, Kasugai Y, Markovic M, Xu C, Fadok JP, Lu T, Poe MM, Xu L, Cook JM, Rudolph U, Sah P, Ferraguti F, Lüthi A. 2015. Regulating anxiety with extrasynaptic inhibition. Nat Neurosci 18:1493–1500. doi:10.1038/nn.4102

Braudeau J, Delatour B, Duchon A, Pereira PL, Dauphinot L, de Chaumont F, Olivo-Marin J-C, Dodd R, Hérault Y, Potier M-C. 2011. Specific targeting of the GABA-A receptor {alpha}5 subtype by a selective inverse agonist restores cognitive deficits in Down syndrome mice. J Psychopharmacol 25:1030–1042. doi:10.1177/0269881111405366

Buzsáki G, Wang X-J. 2012. Mechanisms of Gamma Oscillations. Annu Rev Neurosci 35:203–225. doi:10.1146/annurev-neuro-062111-150444

Cardin JA, Carlésn M, Meletis K, Knoblich U, Zhang F, Deisseroth K, Tsai LH, Moore CI. 2009. Driving fast-spiking cells induces gamma rhythm and controls sensory responses. Nature 459:663–667. doi:10.1038/nature08002

Chang P, Bush D, Schorge S, Good M, Canonica T, Shing N, Noy S, Wiseman FK, Burgess N, Tybulewicz VLJ, Walker MC, Fisher EMC. 2020. Altered Hippocampal-Prefrontal Neural Dynamics in Mouse Models of Down Syndrome. Cell Rep 30:1152–1163.e4. doi:10.1016/j.celrep.2019.12.065

Cho KKA, Hoch R, Lee AT, Patel T, Rubenstein JLR, Sohal VS. 2015. Gamma rhythms link prefrontal interneuron dysfunction with cognitive inflexibility in dlx5/6+/-mice. Neuron 85:1332–1343. doi:10.1016/j.neuron.2015.02.019

Clem RL, Cummings KA. 2020. Prefrontal somatostatin interneurons encode fear memory. Nat Neurosci 23:p61, 14 p. doi:10.1038/s41593-019-0552-7

Davisson MT, Schmidt C, Reeves RH, Irving NG, Akeson EC, Harris BS, Bronson RT. 1993. Segmental trisomy as a mouse model for Down syndrome. Prog Clin Biol Res 384:117–33.

Dawson GR, Maubach KA, Collinson N, Cobain M, Everitt BJ, Macleod AM, Choudhury HI, Mcdonald LM, Pillai G, Rycroft W, Smith AJ, Sternfeld F, Tattersall FD, Wafford KA, Reynolds DS, Seabrook GR, Atack JR. 2006. An Inverse Agonist Selective for 5 Subunit-Containing GABA. J Pharmacol Exp Ther 316:1335–1345. doi:10.1124/jpet.105.092320.amnesic

Deleuze C, Bhumbra GS, Pazienti A, Lourenço J, Mailhes C, Aguirre A, Beato M, Bacci A. 2019. Strong preference for autaptic self-connectivity of neocortical PV interneurons facilitates their tuning to γ-oscillations. PLOS Biol 17:e3000419. doi:10.1371/journal.pbio.3000419

DiGuiseppi C, Hepburn S, Davis JM, Fidler DJ, Hartway S, Lee NR, Miller L, Ruttenber M, Robinson C. 2010. Screening for autism spectrum disorders in children with down syndrome: Population prevalence and screening test characteristics. J Dev Behav Pediatr 31:181–191. doi:10.1097/DBP.0b013e3181d5aa6d

Duchon A, Gruart A, Albac C, Delatour B, Zorrilla de San Martin J, Delgado-García JM, Hérault Y, Potier M-C. 2019. Long-lasting correction of in vivo LTP and cognitive deficits of mice modelling Down syndrome with an a5-selective GABA_A_ inverse agonist. Br J Pharmacol 1–13. doi:10.1111/bph.14903

Erisir A, Lau D, Rudy B, Leonard CS. 1999. Function of Specific K+ Channels in Sustained High-Frequency Firing of Fast-Spiking Neocortical Interneurons. J Neurophysiol 82:2476–2489. doi:10.1152/jn.1999.82.5.2476

Fernandez F, Morishita W, Zuniga E, Nguyen J, Blank M, Malenka RC, Garner CC. 2007. Pharmacotherapy for cognitive impairment in a mouse model of Down syndrome. Nat Neurosci 10:411–413. doi:10.1038/nn1860

Glykys J, Mody I. 2010. Hippocampal Network Hyperactivity After Selective Reduction of Tonic Inhibition in GABA A Receptor ␣ 5 Subunit – Deficient Mice. J Neurophysiol 2796–2807. doi:10.1152/jn.01122.2005

Hannan S, Minere M, Harris J, Izquierdo P, Thomas P, Tench B, Smart TG. 2020. GABA_A_R isoform and subunit structural motifs determine synaptic and extrasynaptic receptor localisation. Neuropharmacology 107540. doi:10.1016/j.neuropharm.2019.02.022

Hausrat TJ, Muhia M, Gerrow K, Thomas P, Hirdes W, Tsukita S, Heisler FF, Herich L, Dubroqua S, Breiden P, Feldon J, Schwarz JR, Yee BK, Smart TG, Triller A, Kneussel M. 2015. Radixin regulates synaptic GABA_A_ receptor density and is essential for reversal learning and short-term memory. Nat Commun 6:6872. doi:10.1038/ncomms7872

Helfrich RF, Knight RT. 2016. Oscillatory Dynamics of Prefrontal Cognitive Control. Trends Cogn Sci. doi:10.1016/j.tics.2016.09.007

Herault Y, Delabar JM, Fisher EMC, Tybulewicz VLJ, Yu E, Brault V. 2017. Rodent models in Down syndrome research: Impact and future opportunities. DMM Dis Model Mech 10:1165–1186. doi:10.1242/dmm.029728

Kaiser T, Ting JT, Monteiro P, Feng G. 2016. Transgenic labeling of parvalbumin-expressing neurons with tdTomato. Neuroscience 321:236–245. doi:10.1016/j.neuroscience.2015.08.036

Kawaguchi Y, Kubota Y. 1998. Neurochemical features and synaptic connections of large physiologically-identified GABAergic cells in the rat frontal cortex. Neuroscience 85:677–701. doi:10.1016/S0306-4522(97)00685-4

Kim D, Jeong H, Lee J, Ghim JW, Her ES, Lee SH, Jung MW. 2016. Distinct Roles of Parvalbumin-and Somatostatin-Expressing Interneurons in Working Memory. Neuron 92:902–915. doi:10.1016/j.neuron.2016.09.023

Kim H, Ährlund-Richter S, Wang X, Deisseroth K, Carlén M. 2016. Prefrontal Parvalbumin Neurons in Control of Attention. Cell 164:208–218. doi:10.1016/j.cell.2015.11.038

Kleschevnikov AM, Belichenko P V., Villar AJ, Epstein CJ, Malenka RC, Mobley WC. 2004. Hippocampal Long-Term Potentiation Suppressed by Increased Inhibition in the Ts65Dn Mouse, a Genetic Model of Down Syndrome. J Neurosci 24:8153–8160. doi:10.1523/JNEUROSCI.1766-04.2004

Larkum M. 2013. A cellular mechanism for cortical associations: An organizing principle for the cerebral cortex. Trends Neurosci. doi:10.1016/j.tins.2012.11.006

Lee NR, Anand P, Will E, Adeyemi EI, Clasen LS, Blumenthal JD, Giedd JN, Daunhauer LA, Fidler DJ, Edgin JO. 2015. Everyday executive functions in Down syndrome from early childhood to young adulthood: evidence for both unique and shared characteristics compared to youth with sex chromosome trisomy (XXX and XXY). Front Behav Neurosci 9:264. doi:10.3389/fnbeh.2015.00264

Lovett-Barron M, Turi GF, Kaifosh P, Lee PH, Bolze F, Sun XH, Nicoud JF, Zemelman B V, Sternson SM, Losonczy A. 2012. Regulation of neuronal input transformations by tunable dendritic inhibition. Nat Neurosci 15:423–430. doi:10.1038/nn.3024

Ma Y. 2006. Distinct Subtypes of Somatostatin-Containing Neocortical Interneurons Revealed in Transgenic Mice. J Neurosci 26:5069–5082. doi:10.1523/JNEUROSCI.0661-06.2006

Martínez-Cué C, Martinez P, Rueda N, Vidal R, Garcia S, Vidal V, Corrales A, Montero JA, Pazos A, Florez J, Gasser R, Thomas AW, Honer M, Knoflach F, Trejo JL, Wettstein JG, Hernandez M-C. 2013. Reducing GABA_A_ 5 Receptor-Mediated Inhibition Rescues Functional and Neuromorphological Deficits in a Mouse Model of Down Syndrome. J Neurosci 33:3953–3966. doi:10.1523/JNEUROSCI.1203-12.2013

Okaty BW, Miller MN, Sugino K, Hempel CM, Nelson SB. 2009. Transcriptional and Electrophysiological Maturation of Neocortical Fast-Spiking GABAergic Interneurons. J Neurosci 29:7040–7052. doi:10.1523/JNEUROSCI.0105-09.2009

Olmos-Serrano JL, Tyler WA, Cabral HJ, Haydar TF. 2016. Longitudinal measures of cognition in the Ts65Dn mouse: Refining windows and defining modalities for therapeutic intervention in Down syndrome. Exp Neurol 279:40–56. doi:10.1016/j.expneurol.2016.02.005

Perrenoud Q, Pennartz CMA, Gentet LJ. 2016. Membrane Potential Dynamics of Spontaneous and Visually Evoked Gamma Activity in V1 of Awake Mice. PLoS Biol 14:e1002383. doi:10.1371/journal.pbio.1002383

Royer S, Zemelman B V., Losonczy A, Kim J, Chance F, Magee JC, Buzsáki G. 2012. Control of timing, rate and bursts of hippocampal place cells by dendritic and somatic inhibition. Nat Neurosci 15:769–775. doi:10.1038/nn.3077

Ruiz-Mejias M, Ciria-Suarez L, Mattia M, Sanchez-Vives M V. 2011. Slow and fast rhythms generated in the cerebral cortex of the anesthetized mouse. J Neurophysiol 106:2910–2921. doi:10.1152/jn.00440.2011

Schulz JM, Knoflach F, Hernandez M-C, Bischofberger J. 2018. Dendrite-targeting interneurons control synaptic NMDA-receptor activation via nonlinear a5-GABA_A_ receptors. Nat Commun 9:3576. doi:10.1038/s41467-018-06004-8

Schulz JM, Knoflach F, Hernandez M, Bischofberger J. 2019. Enhanced dendritic inhibition and impaired NMDAR activation in a mouse model of Down syndrome Section: Neurobiology of Disease Enhanced dendritic inhibition and impaired activation in a mouse model of Down syndrome Department of Biomedicine University of.

Serwanski DR, Miralles CP, Christie SB, Mehta AK, Li X, Blas AL De. 2006. Synaptic and non-synaptic localization of GABA A receptors containing the alpha5 subunit in the rat brain. J Comp Neurol Neurol 499:458–470. doi:10.1002/cne.21115.Synaptic

Silberberg G, Markram H. 2007. Disynaptic Inhibition between Neocortical Pyramidal Cells Mediated by Martinotti Cells. Neuron 53:735–746. doi:10.1016/j.neuron.2007.02.012

Sohal VS, Zhang F, Yizhar O, Deisseroth K. 2009. Parvalbumin neurons and gamma rhythms enhance cortical circuit performance. Nature 459:698–702. doi:10.1038/nature07991

Sternfeld F, Carling RW, Jelley RA, Ladduwahetty T, Merchant KJ, Moore KW, Reeve AJ, Street LJ, O’Connor D, Sohal B, Atack JR, Cook S, Seabrook G, Wafford K, Tattersall FD, Collinson N, Dawson GR, Castro JL, MacLeod AM. 2004. Selective, Orally Active γ-Aminobutyric AcidA a5 Receptor Inverse Agonists as Cognition Enhancers. J Med Chem 47:2176–2179. doi:10.1021/jm031076j

Tremblay R, Lee S, Rudy B. 2016. GABAergic Interneurons in the Neocortex: From Cellular Properties to Circuits. Neuron 91:260–292. doi:10.1016/j.neuron.2016.06.033

Valbuena S, García Á, Mazier W, Paternain A V., Lerma J. 2019. Unbalanced dendritic inhibition of CA1 neurons drives spatial-memory deficits in the Ts2Cje Down syndrome model. Nat Commun 10. doi:10.1038/s41467-019-13004-9

Veit J, Hakim R, Jadi MP, Sejnowski TJ, Adesnik H. 2017. Cortical gamma band synchronization through somatostatin interneurons. Nat Neurosci 20:951–959. doi:10.1038/nn.4562

Wiseman FK, Al-Janabi T, Hardy J, Karmiloff-Smith A, Nizetic D, Tybulewicz VLJ, Fisher EMC, Strydom A. 2015. A genetic cause of Alzheimer disease: Mechanistic insights from Down syndrome. Nat Rev Neurosci 16:564–574. doi:10.1038/nrn3983

Zorrilla de San Martin J, Delabar JM, Bacci A, Potier MC. 2018. GABAergic over-inhibition, a promising hypothesis for cognitive deficits in Down syndrome. Free Radic Biol Med 114:33–39. doi:10.1016/j.freeradbiomed.2017.10.002

Zucca S, D’Urso G, Pasquale V, Vecchia D, Pica G, Bovetti S, Moretti C, Varani S, Molano-Mazón M, Chiappalone M, Panzeri S, Fellin T. 2017. An inhibitory gate for state transition in cortex. Elife 6:1–31. doi:10.7554/elife.26177

